# Dexamethasone Inhibits Cytokine-Induced, DUOX2-Related VEGF-A Expression and DNA damage in Human Pancreatic Cancer Cells and Growth of Pancreatic Cancer Xenografts

**DOI:** 10.1101/2022.02.13.480277

**Authors:** Yongzhong Wu, Mariam M. Konaté, Melinda Hollingshead, Baktiar Karim, Becky Diebold, Jiamo Lu, Smitha Antony, Jennifer L. Meitzler, Agnes Juhasz, Guojian Jiang, Iris Dahan, Krishnendu Roy, James H. Doroshow

## Abstract

Previously, we demonstrated that pro-inflammatory cytokines enhance dual oxidase 2 (DUOX2)-dependent production of reactive oxygen species by human pancreatic ductal carcinoma (PDAC) cells, and that DUOX2 expression is significantly increased in patients with early stages of PDAC. In genetically-engineered mouse models of PDAC, dexamethasone (Dex) decreases formation of pancreatic intraepithelial neoplasia (PanIn) foci as well as PDAC invasiveness. Herein, we report that Dex, in a concentration- and time-dependent fashion, inhibited pro-inflammatory cytokine (IFN-γ/LPS/IL-17A/IL-4)-mediated enhancement of DUOX2 expression in BxPC-3, CFPAC-1, and AsPC-1 human PDAC cell lines, as well as DUOX2–induced DNA damage. The inhibitory effects of Dex were abolished by pre-treatment with the Dex antagonist RU-486. Examination of the human DUOX2 promoter in silico revealed a putative negative glucocorticoid receptor (GR) binding element (IRnGRE). Western analysis, using nuclear extracts from Dex-treated PDAC cells, demonstrated that Dex activated the glucocorticoid receptor in PDAC cell nuclei in the presence of certain co-repressors, such as NCoR-1/2 and histone deacetylases (HDAC1, 2, and 3). Dex produced no anti-proliferative effects on PDAC cells *in vitro*. However, Dex significantly decreased the growth of BxPC-3 xenografts while decreasing inflammatory and immune cell infiltration of the microenvironment, as well as the mRNA expression of DUOX2 and VEGF-A, in BxPC-3 tumors. In contrast, Dex had no effect on the growth of xenografts developed from MIA-PaCa cells that are unresponsive to pro-inflammatory cytokines in culture. In summary, these studies suggest that suppression of inflammation-related DUOX2 expression by Dex could diminish the oxidative milieu supporting PDAC growth and development.

## Introduction

Chronic pancreatitis promotes the initiation and progression of pancreatic cancer, whereas anti-inflammatory interventions that employ either steroid or nonsteroidal anti-inflammatory drugs enhance the repair of inflammation-related tissue injury and decrease subsequent tumorigenesis (1–3). Pro-inflammatory cytokine-, growth factor-, and oncogene-mediated reactive oxygen species (ROS) have been suggested to play an important role in the pathogenesis of exocrine cancers of the pancreas (4–10). ROS production in many epithelial malignancies, including PDAC, may originate from the activity of members of the NADPH oxidase (NOX) gene family (11–14). One of the seven epithelial NADPH oxidases, DUOX2, is highly expressed on the plasma membranes of PDAC cells where it generates hydrogen peroxide; ROS production by DUOX2 also plays an important role in innate immunity at mucosal surfaces in both the respiratory and gastrointestinal tracts and contributes to chronic inflammation-related tissue injury and tumorigenesis (15–18).

In previous studies, we demonstrated that DUOX protein expression was significantly increased in patients with chronic pancreatitis, pancreatic intraepithelial neoplasia (PanIN), early stages of pancreatic cancer, and in human pancreatic cancer xenografts in mice (18, 19). Furthermore, pro-inflammatory cytokines, such as interferon-γ, IL-4, and IL-17A, up-regulate the expression of DUOX2 and its cognate maturation factor DUOXA2 in human pancreatic cancer cells, leading to sustained extracellular H_2_O_2_ accumulation associated with H_2_O_2_-medated DNA double strand scission, and a significant increase in the expression of VEGF and Hif-1α (18–21). Thus, inflammatory cytokine-related, DUOX2-dependent H_2_O_2_ production may contribute to a pro-angiogenic microenvironment rich in ROS that is conducive to the maintenance of genetic heterogeneity; such an environment could enhance the pathologic potential of chronic pancreatic inflammation (8).

Dexamethasone (Dex), a synthetic glucocorticoid (GC) with anti-inflammatory and immunosuppressive properties, is widely used to treat inflammatory, autoimmune, and allergic disorders (22, 23). Its therapeutic effects have been ascribed to its capacity to repress pro-inflammatory cytokine production or cytokine receptors, inflammatory cell infiltration, and/or adhesion molecules activated during the inflammatory response (24, 25). Dex inhibits the invasiveness of human pancreatic cancer cells, in part, by downregulating matrix metalloproteinase expression (26). *In vivo*, Dex alleviates very-early-onset ileocolitis in mice deficient in the antioxidant genes GSH peroxidase 1 and 2 by blocking the upregulation of NOX1, NOX4, and DUOX2 (27). Dex also has demonstrated anticancer activity in PDAC xenograft models (28, 29). Finally, Dex has been reported to inhibit the formation of PanIn foci in genetically-engineered mouse models of pancreatic cancer, as well as the epithelial to mesenchymal transition, and local tumor recurrence and metastasis in other model systems (2,3,30).

The effects of GCs, such as Dex, require the presence of a ligand-activated glucocorticoid receptor (GR) to regulate gene expression. After glucocorticoid binding, activated GRs translocate to the nucleus, homodimerize, and bind to glucocorticoid-responsive elements (GREs) in the 5′-flanking sequences of target gene promoters to alter gene expression positively or negatively (22-25,31,32). The GRE *cis* element to which activated GR binds has been extensively studied in target genes; its consensus sequence is 5’-GGTACAnnnTGTTCT-3’ (23, 33). GREs confer transcriptional activation to agonist-liganded GRs through association with coactivators (e.g., SRC1, TIF2/SRC2 and SRC3) (34). GCs can also induce direct transrepression through binding of GR to inverted repeated negative GC response elements (IRnGRE) with a consensus motif of 5’-CTCC(N)_1-2_GGAG-3’. These elements act on agonist-liganded GR promoting the assembly of cis-repressing complexes containing GR-SMRT/NCoR, and histone deacetylases (HDACs) (35). Alternatively, for some genes that lack functional GRE cis-elements in their promoter regions, ligand-activated GR can regulate gene expression by binding to activated NF-κB or adaptor protein 1 (AP1) to *trans*-repress transcription factor-regulated inflammatory genes (36, 37).

In the current experiments, we found that Dex, in a concentration- and time-dependent fashion, inhibited cytokine (IFN-γ/LPS)-mediated upregulation of DUOX2 and VEGF-A in BxPC-3 and CFPAC-1 human pancreatic cancer cell lines. Dex also inhibited pro-inflammatory cytokine–related DNA damage measured by a significant decrease in γH2AX expression. In BxPC-3 tumor xenografts, Dex also significantly decreased DUOX2 and VEGF-A expression, inflammatory and immune cell infiltration, as well as tumor growth. In contrast, for the MIA-PaCa cell line that is unresponsive to pro-inflammatory cytokines in culture, Dex produced no effect on xenograft growth. Examination of the human DUOX2 promotor in silico revealed a putative negative glucocorticoid receptor binding element (nGRE; 5’-<u>CTCCaGGAG-3’</u>). Western analysis, using nuclear extracts from pancreatic cancer cells treated with Dex, revealed that both activated glucocorticoid receptor and certain co-repressors, such as NCOR-1/2 and histone deacetylases (HDAC1, 2, and 3), exist in human pancreatic cancer cell nuclei. These studies suggest that the inhibitory effects of Dex on PDAC growth *in vivo* may be related to partial suppression of a pro-inflammatory, oxidative tumor milieu.

## Materials and Methods

### Reagents and antibodies

Recombinant human IFN-γ (285-IF0), IL-17A (317-ILB-050), IL-4 (204-IL-050), and IFN-β (8499-IF-010) were purchased from R&D Systems (Minneapolis, MN, USA). All cytokines were dissolved in 0.1% BSA in sterilized PBS. Lipopolysaccharide (LPS, 437620-5MG), dexamethasone (265005-100MG), mifepristone (475838-50MG), and the calcium salt of ionomycin (407952-1MG) were from EMD Millipore (290 Concord Road, Massachusetts 01821, USA). Anti-human DUOX antibody, which reacts with both DUOX1 and DUOX2, was previously developed by Creative Biolabs (Port Jefferson Station, NY, USA) and characterized by our laboratory (14). Anti-Hif-1α (610959) and anti-STAT6 pY641 (611566) were from BD Transduction Laboratories (San Diego, CA, USA). Antibodies against human c-Jun (9165), Jun B (3746), Lamin A/C (2032), ERK1/2 (9102), p-ERK (9101), Hif-1β (3414S), p-histone γH2AX (2577S), p-STAT1_S727_ (9177S), p-STAT1_Y701_ (9167), p-Glucocorticoid Receptor p-GR_S211_ (4161S), p-STAT2_Y690_ (4441S), p-S6_S240/244_ (2215), and p-S6_S235/236_ (4856) were purchased from Cell Signaling Technology (Beverly, MA, USA). Antibodies against human HDAC1 (Ab137652), HDAC2 (Ab137364), HDAC3 (Ab137704), NCoR1 (Ab24552), NCoR2 (Ab24551), and Glucocorticoid Receptor (Ab109022) were from Abcam (Cambridge, MA, USA). STAT1 p84/p91 (sc-346), goat anti-rabbit IgG-HRP (sc-2054), and goat anti-mouse IgG - HRP (sc-2055) were from Santa Cruz Biotechnology. Antibodies used for immunohistochemistry were obtained and used as follows: CD8a (eBioscience #14-0195-82, 1:50), CD45R (BD Biosciences #553086, 1:400), CD45 (BD Biosciences #550539), FoxP3 (eBioscience #14-15773), Microglia Iba-1 (Biocare #CP 290, 1:500), Ly6G/GR1 Granulocyte Marker (Origene #DM3589P, 1:100), and CD3 (BioRad #MCA1477, 1:100 60’). These antibodies recognize murine epitopes. The antibody recognizing Cleaved Caspase-3 was from Cell Signaling Technology (#9661, used 1:100 for 60’); this antibody recognizes both murine and human epitopes. Antibody against Ki67 was obtained from Cell Signaling Technology (#9027, 1:200); this antibody recognizes only human epitopes. All of the human PCR primers: β-actin (Hs01060665_g1), DUOX2 (Hs00204187_m1), DUOXA2 (Hs01595310_g1), STAT1 (Hs00234029_ml), VEGF-A (Hs00173626_ml), and IRF-1 (HS00971965_m1) were purchased from Life Technologies (Carlsbad, CA, USA). Negative Control #1 siRNA (AM4635), human DUOX2 siRNA ‘A’ (S27012), and human DUOX1 siRNA (S28797) were obtained from Applied Biosystems (Carlsbad, CA, USA). DUOX2 siRNA ‘B’ (Cat# J-008324-05-0020) was from Horizon Discovery. Diphenylene iodonium (DPI) (D2926), n-acetyl-L-cysteine (NAC) (A7250), and PEG-catalase (C4963) were from Sigma-Aldrich (St. Louis, MO, USA).

### Cell culture

The human pancreatic cancer cell lines BxPC-3 (CRL-1687), AsPC-1 (CRL-1682), MIA-PaCa (CRL-1420), and CFPAC-1 (CRL-1918) were obtained from the American Type Culture Collection (ATCC) (Manassas, VA, USA). Both BxPC-3 and AsPC-1 cells were cultured in RPMI 1640 medium (SH30255.01) (HyClone, Logan, UT, USA) supplemented with 1.0% sodium pyruvate and 10% FBS (100-106; Gemini Bio Products, Sacramento, CA, USA). For MIA-PaCa cells, the complete medium was Dulbecco’s Modified Eagle’s Medium (DMEM, ATCC# 30-2002) plus 10% FBS and 2.5% horse serum. CFPAC-1 cells were cultured in Iscove’s Modified Dulbecco’s medium with 10% FBS. The identity of each cell line was confirmed by the Genetic Resources Core Facility of Johns Hopkins University (Baltimore, MD, USA). To establish starvation conditions before each experiment, cells were cultured overnight in the same medium without FBS. Starvation conditions were used because DUOX2 induction by different cytokines is stronger after serum starvation, as noted previously. In all cases, cells were cultured in a humidified incubator at 37°C in an atmosphere of 5% CO_2_ in air.

### Transfection with siRNA and cell treatment for RNA and protein analysis

Lonza Kit V (VCA-1003) was used to introduce siRNA into BxPC-3 cells with the Lonza Nucleofector 2b device (Cat# AAB-1001) following the manufacturer’s protocol. Cells (1×10^6^) were resuspended in the provided Kit V electroporation solution, and control or gene specific siRNAs with a 20 nM final concentration were added to the cell suspension before transferring to a clean electroporation cuvette. After the electric charge was applied, cells were transferred with a transfer pipette to culture dishes containing full serum and media. The cell-type specific program was used for the electroporation procedure. Lipofectamine RNAIMAX reagent (Thermo Fisher Scientific, Cat# 13778075) was used to transfect siRNA into CFPAC-1 cells following the company’s instructions both for control and gene specific siRNAs with a final siRNA concentration of 20 nM. For RNA and Amplex Red assays, 1×10^6^ cells were seeded into 60 mm dishes with 4 ml of complete medium; for protein assays, 4 ×10^6^ cells were plated into 100 mm dishes with 10 ml of complete medium.

### Mice

All mice (female athymic nu/nu NCr) used in the study were bred in-house from production stock originating from the NCI-Frederick Animal Production Program, Frederick, MD. National Cancer Institute Frederick is accredited by the American Association for the Accreditation of Laboratory Animal Care International and follows the Public Health Service Policy for the Care and Use of Laboratory Animals. Animal care was provided in accordance with the procedures outlined in the Guide for the Care and Use of Laboratory Animals. All studies were conducted under an approved Institutional Animal Care and Use Committee protocol.

### Quantitative real-time RT-PCR

Total RNA was extracted from cells using the RNeasy Mini Kit (74104; Qiagen, Valencia, CA, USA) following the manufacturer’s protocol. Two micrograms of total RNA were then used for cDNA synthesis, along with Super Script II Reverse Transcriptase (18080–044) and random primers (48190–011) (both from Life Technologies), in a 20 µl reaction. The reaction conditions were: 25°C for 10 min, 42°C for 50 min, 75°C for 10 min, and 4°C indefinitely. Real-time quantitative PCR products were diluted to 100 µl with diethylpyrocarbonate-treated H_2_O, and real-time quantitative PCR was conducted in 384-well plates in a 20 µl volume consisting of 2 µl of diluted cDNA, 1 µl of the appropriate primers, 7 µl of H_2_O, and 10 µl of TaqMan Universal PCR Master Mix (4364340; Life Technologies). The PCR reaction was performed using the default cycling conditions (50°C for 2 min, 95°C for 10 min, and 40 cycles of 95°C for 15 sec and 60°C for 10 min) with the ABI Prism 7900HT Sequence Detection System (Applied Biosystems). Triplicate samples were used for real-time quantitative PCR, and the mean values were calculated. The data in all figures represent three independent experiments. Relative gene expression was calculated from the ratio of the target gene expression to the internal reference gene (β-actin) expression based on the Ct values.

### Western analysis

To prepare extracts, cell pellets were lysed in 1X RIPA Lysis Buffer (20-188; EMD Millipore) supplemented with phosphatase inhibitor (04-906-837001) and protease inhibitor (11-836-153001s) (both from Roche, Mannheim, Germany) to generate whole-cell extracts (WCEs). Nuclear extracts were prepared using the NE-PER Nuclear and Cytoplasmic Extraction Kit (78833; Thermo Scientific, Rockford, IL, USA). The protein concentrations of both types of extracts were measured using the BCA Protein Assay Kit (23227; Pierce, Rockford, IL, USA). The lysates were then combined with an equal volume of 2X SDS Protein Gel Loading buffer (351-082-661; Quality Biological, Gaithersburg, MD, USA). Next, unless otherwise described, 50 μg of WCE or 20 μg of nuclear extract was electrophoretically separated on a 4-20% Tris-glycine gel (EC6028; Life Technologies) and transferred to nitrocellulose membranes using an iBlot 2 Transfer Stack (IB 3010-01; Life Technologies). The membranes were blocked in 5% nonfat milk in 1X TBST (TBS with 0.1% Tween 20) buffer for 1 h at room temperature and then incubated overnight with the indicated primary antibodies in TBST. After three washes with TBST, the membranes were incubated with the appropriate HRP-conjugated secondary antibodies for 1 h at room temperature on a shaker. SuperSignal West Pico Luminol/Enhancer Solution (1856136; Thermo Scientific) was then applied to visualize the proteins of interest.

### Amplex Red^®^ assay to detect extracellular H_2_O_2_

The Amplex Red^®^ Hydrogen Peroxide/Peroxidase Assay Kit (A22188; Life Technologies) was employed to detect extracellular H_2_O_2_ release. Cells were washed twice with 1X PBS, trypsinized, and counted. Next, for BxPC-3 or CFPAC-1 cells, 20 µl of cell suspension containing 2 × 10^4^ live cells in 1X Krebs-Ringer phosphate glucose (KRPG) buffer was mixed with 100 µl of a solution containing 50 µM Amplex Red^®^ and 0.1 U/ml HRP in KRPG buffer with 1 µM ionomycin and incubated at 37°C for the indicated times. The fluorescence of the oxidized 10-acetyl-3,7-dihydroxyphenoxazine was then measured at excitation and emission wavelengths of 530 nm and 590 nm, respectively, using a Spectra Max Multi-Mode Microplate Reader (Molecular Devices, Sunnyvale, CA, USA), and the amount of extracellular H_2_O_2_ was determined using a standard curve from 0-2 µM H_2_O_2_. Each value in the figures is the mean value for triplicate or quadruplicate samples.

### MTT assay for cell proliferation

BxPC-3 or MIA-PaCa cells were seeded in 96-well plates at a density of 1 × 10^4^ cells/well in 200 µl of complete medium overnight. The next day, medium was changed to add solvent (DMSO) or 1 µM of dexamethasone in complete medium. The final DMSO concentration to which the cells were exposed was 0.1% vol/vol. On days 1, 2, and 3 after treatment, cell proliferation was evaluated by the MTT method. In short, the culture medium was removed, and 100 μl of MTT (0.5 mg/ml) was added to each well. The plates were then incubated at 37°C for 45 min. The MTT was removed, and 100 μl of MTT solvent was added to each well. The plates were shaken for 10 min and read at 570 nm with the Spectra Max M5 reader.

### RNA extraction from tumor xenografts

In addition to the methods described above for cell lines, a Polytron System PT 1200E homogenizer equipped with a 5–12-mm-diameter dispersing head (KINEMATICA, Lucerne, Switzerland) was used to homogenize tumor tissue mixed with a suitable amount of RLT buffer for RNA extraction. A 350-μl volume of tumor homogenate was processed directly for isolation of RNA.

### Tumor growth, treatment with dexamethasone *in vivo*, and tumor sampling

Human pancreatic tumor cells for mouse inoculation were used at *in vitro* passages four to six from cryopreserved cell stocks. BxPC-3 or MIA-PaCa tumor cells (1 × 10^7^ cells/0.1 ml/injection) were inoculated subcutaneously into mice (n = 199 mice/tumor cell line). This provided a total of 104 potential tumor samples. On the day of tumor staging, the selected mice were randomized into one of 4 groups using the randomization program provided in Study Director software (StudyLog, South San Francisco, CA). Mice were placed into one of 3 groups: (1) vehicle (0.45% saline) controls (n=32); (2) dexamethasone 12.5 mg/kg every 12 h subcutaneously for 22 doses (n=32); or (3) dexamethasone 6.25 mg/kg every 12 h subcutaneously for 22 doses (n=32). A fourth group (n=8) was randomized into the study to serve as untreated day 0 controls, and these tumors were collected on the first day of the study. Dexamethasone (NDS 63323-165-30) was obtained from the Department of Pharmacy at the NIH Clinical Center, Bethesda, MD. Treatment was initiated and the tumors were monitored for growth using caliper measurements as described previously (38). Over the course of the study, 8 mice were randomly selected from each group for tumor collection 4 h after the last dose of dexamethasone. Tumors were resected and cut into quadrants. One quadrant was placed into 10% neutral buffered formalin; one quadrant into RNALater; and the remaining 2 quadrants were flash frozen. Flash freezing was accomplished by placing the tumor quadrants into pre-chilled (liquid nitrogen [LN2]) cryovials, followed by immediate submersion into LN2. All frozen samples were transferred to a −70°C freezer, and the RNALater samples were refrigerated for holding prior to processing.

### Effect of dexamethasone on immune and inflammatory cell infiltration and markers of proliferation in BxPC-3 xenografts

Evaluation of immune and inflammatory cell infiltration and analysis of proliferation markers were performed on tumors from BxPC-3-bearing mice that had received saline or Dex (12.5 mg/kg, every 12 h for 22 doses); tumors were removed as previously described 4 h after the last dose of Dex; eight mice per group were analyzed. Immunohistochemistry (IHC) was performed using automated staining on a LeicaBiosystems BondRX instrument with the following conditions: Epitope Retrieval 1 (Citrate) 20’ for Cleaved Caspase-3, CD8a, CD45R, Microglia Iba-1, Ly6G/GR1 Granulocyte Marker, CD3, and Ki67. The Bond Polymer Refine Detection Kit (LeicaBiosystems #DS9800) with omission of the PostPrimary Reagent was used, and an anti-rat secondary antibody was included for CD8a, CD45R, Ly6G/GR1, and CD3. Manual staining was performed for CD45 and FoxP3. Following antigen retrieval with citrate buffer, slides were incubated with CD45 (1:100 overnight, room temperature) and FoxP3 (1:100, 1 h), followed by biotinylated anti-rat secondary antibody, ABC Elite (Vector Labs), DAB, and hematoxylin counterstaining. Isotype control reagents were used in place of the primary antibodies for the negative controls. Slides were also stained with H&E following standard protocols. H&E and IHC slides were scanned at 20X using an Aperio AT2 scanner (Leica Biosystems, Buffalo Grove, IL) into whole slide digital images. All image analysis was performed using Halo imaging analysis software (v3.3.2541.300; Indica Labs, Corrales, NM), and image annotations were performed by one pathologist (B.O.K). Areas of artifact such as folds and tears were excluded from analysis.

### Statistical analysis

Statistical differences between the mean values of samples were assessed using the two-tailed Student’s t test; statistical significance was defined as \**P* < 0.05, \*\**P* < 0.01, and \*\*\**P* < 0.001. The two-tailed Student’s t test was also used to assess differences in tumor volumes between control and dexamethasone treated groups.

## Results

### Dex inhibits cytokine-stimulated DUOX2 mRNA and protein expression in BxPC-3 human pancreatic cancer cells

The pro-inflammatory cytokine IFN-γ upregulates DUOX2 mRNA and protein expression in the human pancreatic cancer cell line BxPC-3 (20, 21). We used real time PCR to evaluate the effect of Dex on IFN-γ-induced DUOX2 mRNA expression in these cells. As demonstrated in **Fig. 1A**, consistent with our previous results, IFN-γ significantly increased the mRNA expression of DUOX2 and its cognate maturation factor DUOXA2 in BxPC-3 cells. IFN-γ-enhanced DUOX2 and DUOXA2 levels were significantly inhibited by Dex at a concentration as low as 1 nM (*P* < 0.01 vs. IFN-γ-treated cells not exposed to Dex). Inhibition of DUOX2/DUOXA2 expression by Dex was concentration dependent from 1 to 100 nM. Temporal analysis (**Fig. 1B**) revealed that pretreatment with 0.1 µM Dex for 30 min (prior to a 24 h exposure to IFN-γ) resulted in a significant decrease in the level of DUOX2 mRNA (*P* < 0.001 vs. 0 h pretreatment); Dex pretreatment decreased IFN-γ-induced DUOX2 mRNA expression in a time-dependent fashion from 3 to 24 h.

**Figure 1.**
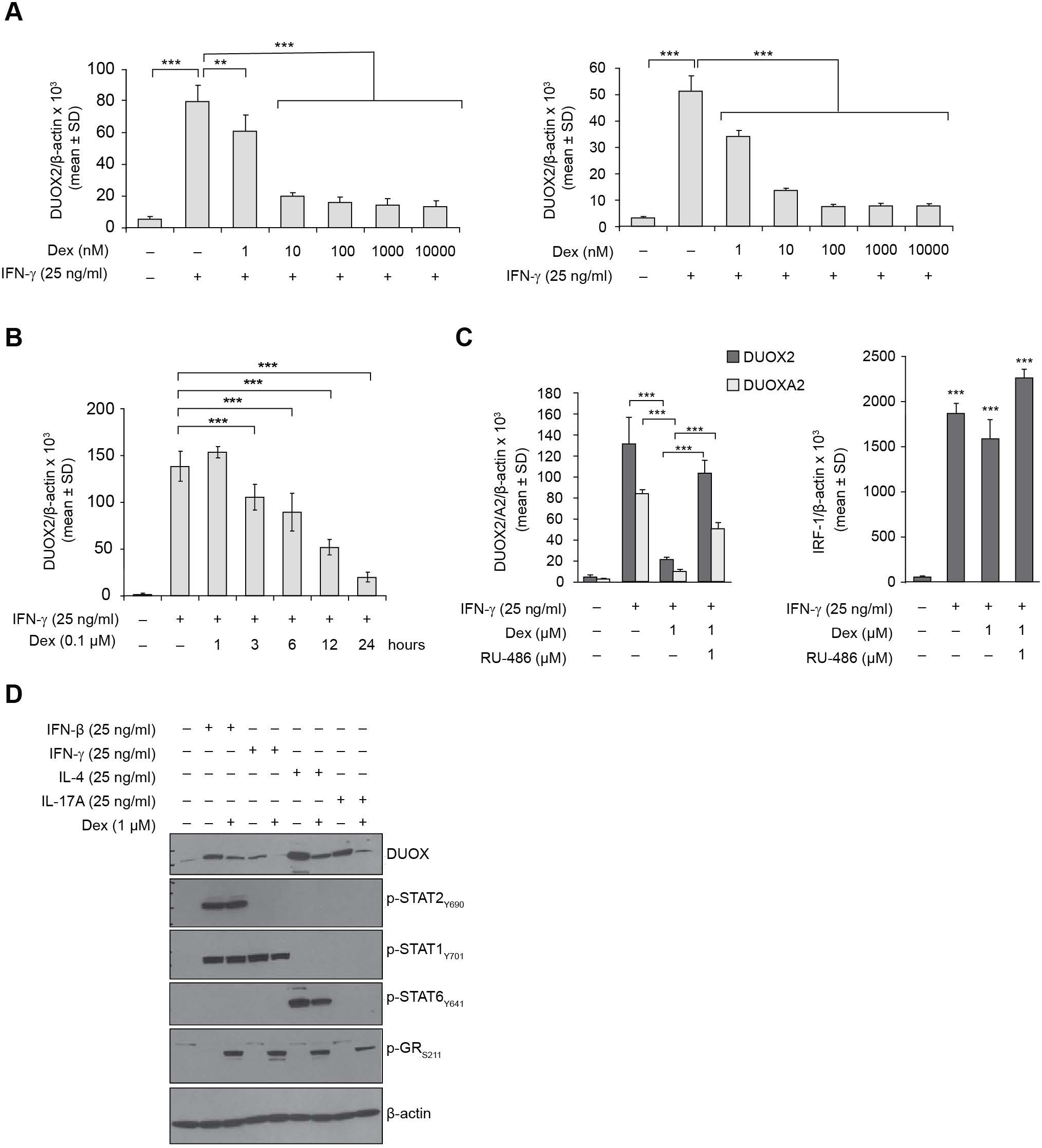
Dexamethasone (Dex) inhibits cytokine-stimulated DUOX2 mRNA and protein expression in BxPC-3 human pancreatic cancer cells. **A,** Dex inhibits DUOX2/DUOXA2 mRNA expression in BxPC-3 cells in a concentration–dependent manner. BxPC-3 cells grown in serum free medium were pretreated with different concentrations of Dex for 30 min and then exposed to IFN-γ (25 ng/ml) for 24 h; cells were collected and subjected to RNA extraction and real time PCR for DUOX2 and DUOXA2 expression relative to β-actin. Error bars represent ± SD; data are from triplicate samples with 9 readings. \*\**P* < 0.01 and \*\*\**P* < 0.001 vs. cells treated with IFN-γ without Dex. **B,** Time course for Dex inhibition of IFN-γ-induced DUOX2 RNA expression in BxPC-3 cells. Quantitative PCR analysis of DUOX2 mRNA expression normalized to β-actin in BxPC-3 cells following exposure to IFN-γ (25 ng/ml) for 24 h with Dex (0.1 µM) pretreatment at different times as indicated in the figure. \*\*\**P* < 0.001 vs. cells treated with IFN-γ but without inhibitor. Results represent triplicate experiments. **C,** Mifepristone (RU-486), a Dex antagonist, relieves Dex-related inhibition of IFN-γ-enhanced DUOX2 RNA expression in BxPC-3 cells. BxPC-3 cells grown in serum free medium were first treated with either 1 µM Dex or 1 µM Dex plus 1µM RU486 for 30 min and then incubated with IFN-γ (25 ng/ml) for 24 h; cells were then collected and RNA extracted for quantitative PCR analysis of (left panel) DUOX2 (dark bar) or DUOXA2 (gray bar) and (right panel) IRF-1 mRNA expression relative to β-actin, *n* = 3. \*\*\**P* < 0.001 for cells treated with IFN-γ alone vs. solvent; \*\*\**P* < 0.001 Dex plus IFN-γ (25 ng/ml) for 24 h vs. cells treated with IFN-γ (25ng/ml) for 24 h alone. \*\*\**P* < 0.001 for cells pretreated with RU-486 plus Dex and then exposed to IFN-γ for 24 h compared to Dex alone followed by IFN-γ (25 ng/ml) for 24 h. **D**, Dex suppresses proinflammatory cytokine-induced DUOX2 protein expression in BxPC-3 cells. Western analysis of whole cell extracts (WCEs) (50 µg) from BxPC-3 cells treated with various cytokines in serum-free medium for 24 h with or without 1 µM of Dex pretreatment for 30 min. β-actin served as a loading control. The data are representative of at least three independent experiments.

Ligand-activated GR is a prerequisite for the regulation of target genes by glucocorticoids (23, 24). To establish whether the inhibition of IFN-γ-mediated upregulation of DUOX2 expression by Dex required the presence of agonist-activated GR, BxPC-3 cells were pretreated for 30min with either solvent (DMSO), or Dex (1 µM), or Dex (1 µM) plus RU-486 (1 µM), a GR antagonist, followed by IFN-γ (25 ng/ml) for 24 h. As demonstrated in the left panel of **Fig. 1C**, IFN-γ-enhanced DUOX2 and DUOXA2 mRNA expression were each significantly decreased by Dex (*P* < 0.001 vs. IFN-γ treatment without Dex). Pretreatment with RU-486, on the other hand, relieved the Dex-mediated inhibition of DUOX2 and DUOXA2 expression by nearly 80%. However, as demonstrated in **Fig. 1C** (right panel), neither Dex nor RU-486 produced an inhibitory effect on IFN-γ-induced IRF-1 expression. This finding suggests that ligand-activated GR is necessary for Dex-mediated inhibition of DUOX2 expression in BxPC-3 cells.

Next, we determined whether the Dex-mediated inhibition of cytokine-enhanced DUOX2 mRNA levels decreased DUOX protein expression. BxPC-3 cells propagated in serum-free medium were first treated with either solvent or 1 µM Dex for 30 min, and then exposed to a variety of pro-inflammatory cytokines (25 ng/ml) for 24 h; cell lysates were subjected to Western analysis. Dex treatment resulted in GR activation, indicated by phosphorylation of Serine 211 on the GR (**Fig. 1D**); Dex decreased cytokine-enhanced DUOX protein expression for each cytokine studied, albeit to varying degrees. For example, Dex almost completely abolished IFN-γ- and IL-17A-enhanced DUOX protein expression; this effect was observed to a lesser degree for IFN-β-and IL-4-stimulated DUOX. Phosphorylation of relevant downstream signaling proteins was observed for each cytokine: STAT2_Y690_ and STAT1_Y701_ for IFN-β; STAT1_Y701_ for IFN-γ; and STAT6_Y641_ for IL-4. Taken together, these results demonstrate that in a GR-dependent manner Dex broadly inhibits pro-inflammatory cytokine-enhanced DUOX protein expression in BxPC-3 human pancreatic cancer cells.

### Dex decreases IFN-γ- and LPS-enhanced DUOX1/2 and VEGF-A mRNA and Hif-1α and c-Jun protein expression in human pancreatic cancer cells

In prior investigations, we demonstrated that IFN-γ and LPS increase DUOX2 and VEGF-A expression concomitantly in human pancreatic cancer cells in a ROS-dependent manner (19). Because Dex can inhibit IFN-γ-enhanced DUOX2/DUOXA2 expression in BxPC-3 cells, we examined whether IFN-γ-enhanced VEGF-A expression was also inhibited in the same cells. As shown in the left and middle panels of **Fig. 2A**, IFN-γ significantly increased both DUOX2 (and to a lesser extent DUOX1) and VEGF-A mRNA expression in BxPC-3 cells (*P* < 0.001 vs. solvent control for DUOX2 and VEGF-A; *P* < 0.05 for DUOX1); IFN-γ-enhanced DUOX2 and VEGF-A mRNA expression were both significantly inhibited by Dex pretreatment (*P* < 0.001 vs. IFN-γ treatment without Dex). In the right panel of **Fig. 2A**, Western analysis using nuclear extracts from BxPC-3 cells treated with solvent or Dex followed by IFN-γ exposure for 24 h demonstrated that Dex treatment was followed by translocation of phosphorylated GR to the nucleus without influencing total GR nuclear distribution. Furthermore, Dex inhibited IFN-γ-enhanced Hif-1α and c-Jun nuclear expression (compare lane 3 for IFN-γ plus solvent to lane 4 for IFN-γ plus Dex).

**Figure 2.**
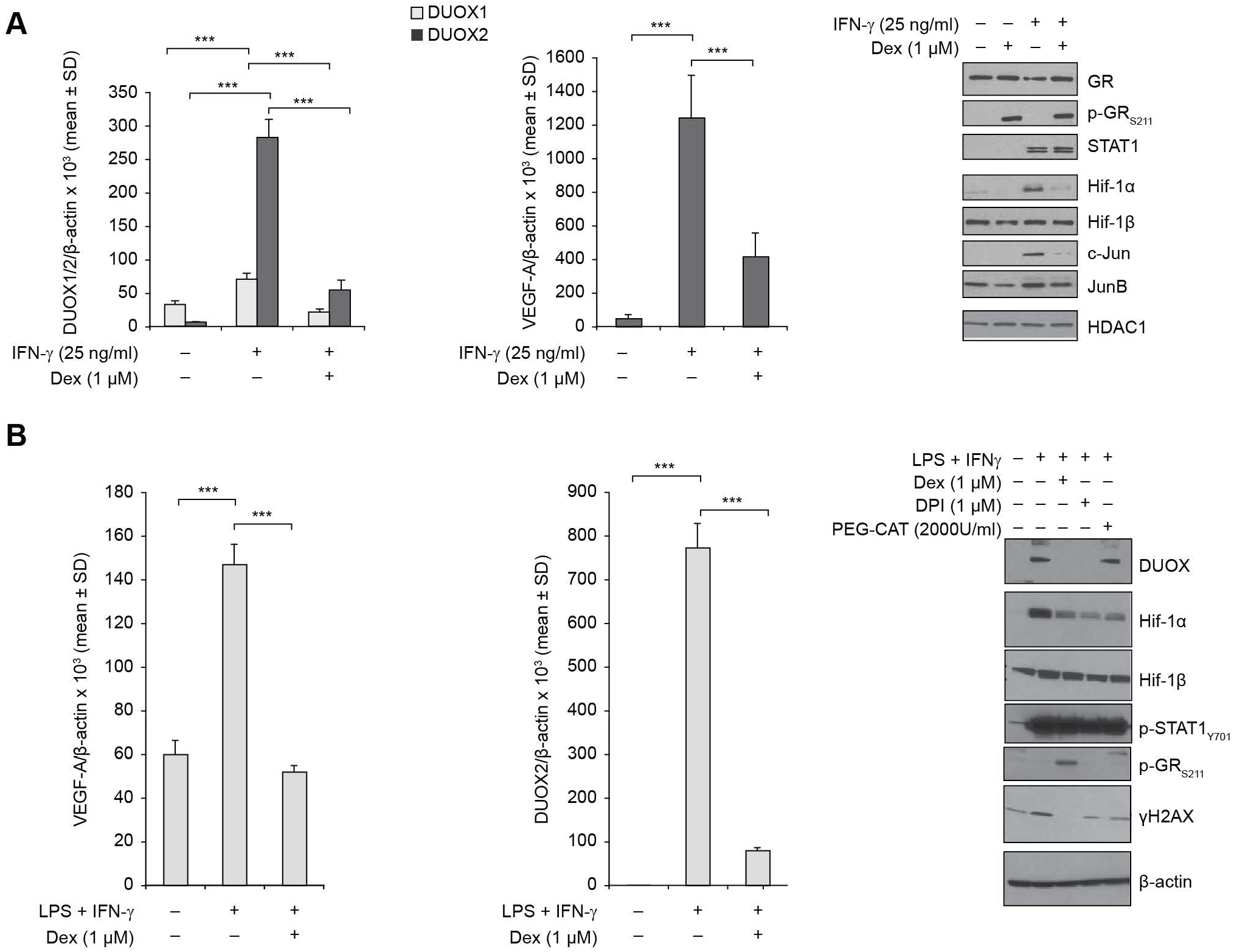
Dexamethasone decreases IFN-γ-induced DUOX2 and VEGF-A mRNA expression and Hif-1α and c-Jun protein expression in human pancreatic cancer cells. **A,** Dex decreases IFN-γ induced DUOX2 and VEGF-A RNA expression in BxPC-3 cells. Real time PCR was used to measure relative DUOX1 (gray) and DUOX2 (black) mRNA expression in the left panel, and VEGF-A mRNA expression in the middle panel normalized to β-actin in BxPC-3 cells treated with IFN-γ (25 ng/ml) for 24 h with or without 1 µM of Dex as a 30 min pretreatment. Dex significantly decreased both DUOX1 (\*\*\**P* < 0.001) and DUOX2 (\*\*\**P* < 0.001) expression vs. solvent treated control cells exposed to IFN-γ (25 ng/ml). Dex also significantly diminished the IFN-γ-mediated increase in VEGF-A expression (\*\*\**P* < 0.001 vs. solvent treated controls). Data represent a minimum of three separate experiments. The Western analysis shown in the right panel employed 20 µg of nuclear extract from BxPC-3 cells treated for 24 h with solvent (lane1), 1 µM Dex alone for 24 h (lane 2), 25 ng/ml of IFN-γ alone for 24 h (lane 3) or IFN-γ for 24 h plus 1µM of Dex as a 30 min pre-treatment. HDAC1 expression serves as the loading control for this experiment that was conducted in triplicate. **B,** Dex suppresses IFN-γ- and LPS-enhanced VEGF-A, DUOX2, and Hif-1α expression in CFPAC-1 cells. Quantitative PCR assay of relative VEGF-A (left panel) or DUOX2 (middle panel) mRNA expression normalized to β-actin in CFPAC-1 cells treated with 1 µg of LPS plus IFN-γ (25 ng/ml) for 24 h with or without 1 µM Dex for a 30 min pretreatment. \*\*\**P* < 0.001 for solvent treated cells vs. LPS plus IFN-γ for 24 h, or LPS plus IFN-γ treatment for 24 h alone vs. LPS plus IFN-γ (25 ng/ml) for 24 h with 1 µM Dex as a 30 min pretreatment. The right panel demonstrates a Western blot using 50 µg whole cell extract from CFPAC-1 cells treated for 24 h with solvent, LPS plus IFN-γ alone or with 1 µM of Dex, 1 µM of DPI or 2000 U/ml PEG-catalase for 30 min as a pretreatment. Results are representative of three experiments.

To explore the effect of Dex on the expression of DUOX2 and the pro-angiogenic factors Hif-1α and VEGF-A in a different model of human pancreatic cancer, CFPAC-1 cells were first treated with solvent or Dex (1 µM) for 30 min and then for 24 h with the combination of IFN-γ (25 ng/ml) and LPS (1 µg/ml). VEGF-A and DUOX2 mRNA expression were investigated, as demonstrated in **Fig. 2B**, left and middle panels. LPS plus IFN-γ strongly increased VEGF-A (*P* < 0.001 vs. solvent treatment) and DUOX2 (*P* < 0.001 vs. solvent treatment) expression. Cytokine-enhanced DUOX2 and VEGF-A levels were significantly inhibited by Dex pretreatment (*P* < 0.001 vs. LPS plus IFN-γ exposure without inhibitor). Western analysis (**Fig. 2B**, right panel, lane 3) demonstrated that Dex enhanced the phosphorylation of GR at serine 211, in a fashion similar to that observed for BxPC-3 cells. Dex strongly inhibited LPS plus IFN-γ (compare lane 2 with lane 3)-enhanced DUOX expression in CFPAC-1 cells. The levels of Hif-1α and the DNA damage marker γ-H2AX were also inhibited by Dex. Furthermore, the NADPH oxidase/dehydrogenase enzyme inhibitor DPI (lane 4) and the H_2_ O_2_ detoxifying enzyme PEG-catalase (lane 5) decreased ROS-mediated Hif-1α and γ-H2AX upregulation by LPS and IFN-γ co-treatment in CFPAC-1 cells.

To provide additional context for these findings, we compared the effects of IFN-γ and Dex on DUOX2 expression and GC signaling in BxPC-3 cells to those in the MIA-PaCa pancreatic cancer cell line that is known to lack measurable constitutive DUOX2 expression (14). As shown in the left panel of Supplementary Fig. S1 and reported above, IFN-γ enhanced both DUOX2 and VEGF-A mRNA expression in BxPC-3 cells; and DUOX2 and VEGF-A expression levels were significantly decreased by pretreatment with Dex. On the other hand, in contemporaneous experiments, neither DUOX2 nor VEGF-A mRNA expression was altered by IFN-γ treatment in MIA-PaCa cells. Hence, as expected, Dex also did not inhibit either DUOX2 or VEGF-A expression in this cell line. Western analysis (right panel) confirmed that IFN-γ did not increase DUOX or Hif-1α protein expression in MIA-PaCa cells, although in this line IFN-γ can activate STAT1 as indicated by STAT1_Y701_ phosphorylation, and Dex treatment leads to GR phosphorylation. These results support our previous finding that increased normoxic Hif-1α expression in BxPC-3 and CFPAC-1 cells is related to DUOX2-mediated ROS production in these models.

To characterize the role of DUOX2-related ROS production on VEGF-A expression in PDAC cells, we assessed the consequences of treatment with the flavin dehydrogenase and NOX inhibitor diphenylene iodonium (DPI) and the multifunctional reduced thiol N-Acetyl-L-Cysteine (NAC) in BxPC-3 cells. As shown in the left panel of Supplementary Fig. S2, NAC and DPI treatment produced a significant decrease in VEGF-A expression in cytokine-treated cells (*P* < 0.001 vs. IFN-γ-treated cells not exposed to inhibitors) without influencing DUOX2 mRNA expression (Supplementary Fig. S2, right panel). In addition to inhibiting the enzymatic activity of DUOX2/DUOXA2 with DPI, to confirm the role of DUOX2-related ROS in the cytokine-mediated upregulation of proangiogenic molecules, we knocked down the expression of DUOXA2 (required for the production of H_2_O_2_) using RNA interference. These studies confirmed that DUOXA2 siRNA not only decreased IFN-γ-enhanced DUOXA2 expression in BxPC-3 cells by more than 90% (*P* < 0.001 vs. control; Supplementary Fig. S3, left panel), but also significantly diminished IFN-γ-mediated VEGF-A upregulation (*P* < 0.001 vs. control; Supplementary Fig. S3, right panel). Together, our data support the conclusion that the upregulation of VEGF-A by IFN-γ in BxPC-3 cells depends on DUOX2-derived ROS.

### Dex inhibits cytokine-induced signal transduction, DNA damage, and H_2_O_2_ production in human pancreatic cancer cells that is mediated by DUOX2

Inflammatory cytokine-related, DUOX2-dependent H_2_O_2_ production can produce a DNA damage response in human pancreatic cancer cells and has been suggested to play an important role in chronic inflammation-associated pancreatic cancer initiation and progression (21). Dex has also been shown to enhance the repair of inflammation-related tissue injury and decrease subsequent tumorigenesis (3). Hence, we evaluated whether Dex could attenuate cytokine-enhanced, DUOX2-associated DNA damage in human pancreatic cancer cells. To this end, the effect of Dex and its antagonist RU486 on IFN-γ-enhanced DUOX2 expression and subsequent activation of relevant signaling pathways was explored in BxPC-3 cells. As shown in **Fig. 3A**, IFN-γ activated STAT and MAPK pathways in BxPC-3 cells, demonstrated by phosphorylation of STAT1 and ERK, respectively. These results are supported at the mRNA level by the inhibition of IFN-γ-related upregulation of VEGF-A following exposure to the MEK inhibitor U01226 (Supplementary Fig. S4). DUOX, c-Jun, Jun B, and γH2AX expression were all increased by IFN-γ and inhibited by Dex. The GR antagonist RU-486 blocked the effect of Dex on IFN-γ-induced ERK activation, and DUOX, c-Jun, JunB, and γH2AX expression. Furthermore, Dex enhanced, and RU486 blocked, the phosphorylation of GR at Ser 211. These data suggest that Dex inhibits IFN-γ-mediated-DUOX2 induction and the subsequent DNA damage response in BxPC-3 cells in a GR dependent manner.

**Figure 3.**
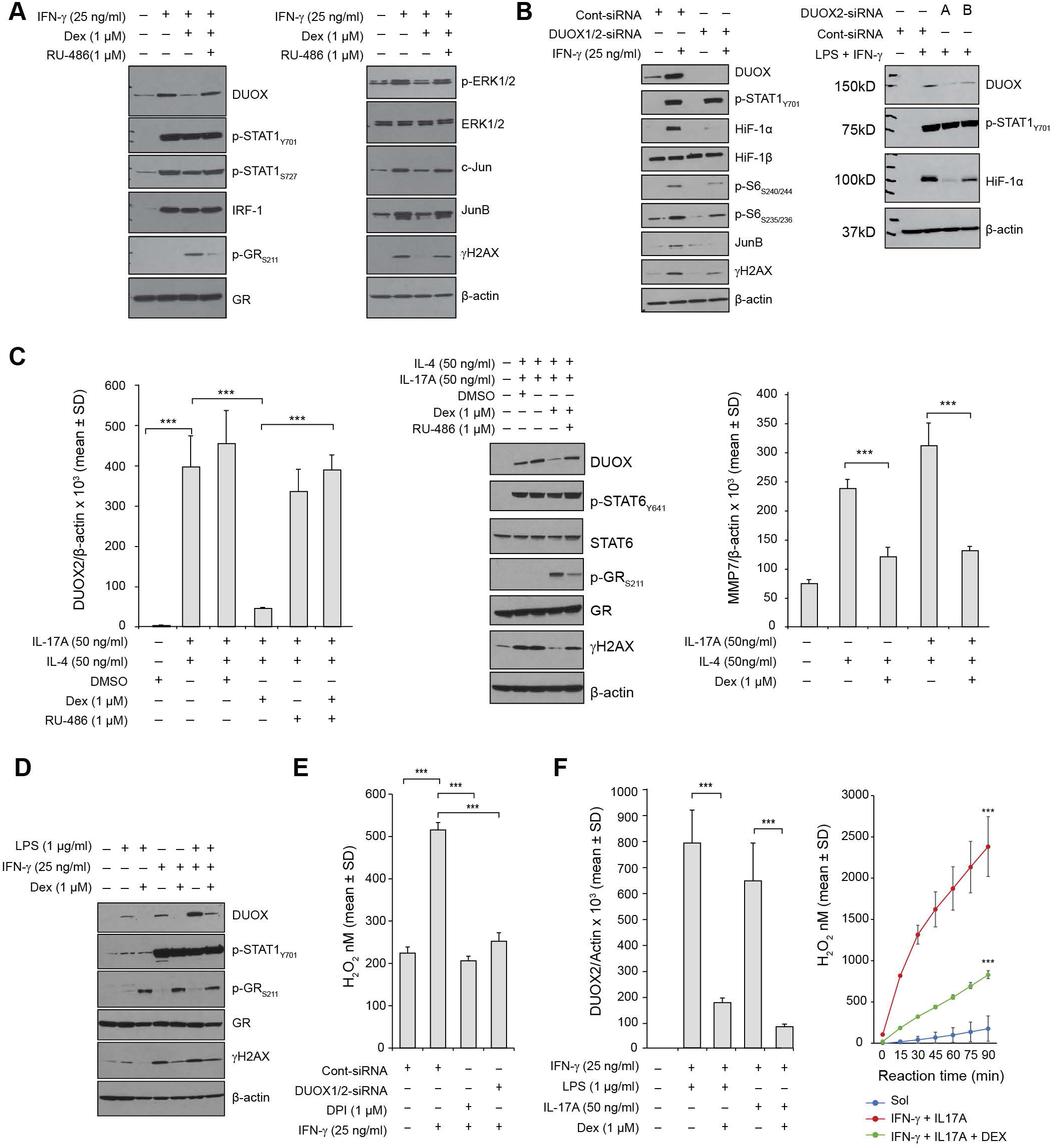
Effect of dexamethasone on cytokine-stimulated signal transduction and DNA damage in human pancreatic cancer cells. **A,** Dex decreases IFN-γ-enhanced DUOX2 protein expression and DNA double strand breaks in BxPC-3 cells. BxPC-3 cells were treated with IFN-γ or solvent for 24 h; Dex alone or Dex plus RU-486 were administered 30 min prior to the initiation of IFN-γ exposure to evaluate their effects on DUOX protein expression and the DNA damage response indicated by the presence of an enhanced phosphorylated histone γH2AX signal. Western analysis was performed with 50 μg of whole cell lysate using the specific antibodies as indicated. The results shown are typical of three identical experiments. **B,** Effect of silencing DUOX1/2 on IFN-γ-induced DUOX protein expression and DNA damage response in BxPC-3 cells (left panel). Control or a combination of both DUOX1 and DUOX2 siRNAs were transiently transfected into BxPC-3 cells; this combination was examined here because IFN-γ produces a small increase in DUOX1 expression as shown in Fig. 2A. At 24 h after transfection, cells were incubated in serum-free medium with or without IFN-γ (25 ng/ml) for a subsequent 12 h, and 50 μg of whole cell extract was subjected to Western analysis, using specific antibodies as indicated. The data are representative of at least three independent experiments. In the right panel, a related experiment in CFPAC-1 cells was conducted using two independent DUOX2 siRNAs and exposure to LPS (1 µg/ml) and IFN-γ (25 ng/ml). CFPAC-1 cells were transfected with either a control siRNA or two different human DUOX2 siRNAs. 24 h following transfection the cells were treated with solvent or LPS and IFN-γ for another 24 h. Western blotting was performed using 50 µg of whole cell lysate. **C,** Dex suppresses IL-4 plus IL-17A-induced DUOX2 and MMP7 expression, as well as DNA damage in AsPC-1 human PDAC cells. Left panel, starved AsPC-1 cells were treated with DMSO or IL-4 (50 ng/ml) plus IL-17A (50 ng/ml) for 24 h with or without either Dex alone or Dex plus RU-486 pretreatment for 30 min; 2 μg of total RNA was subjected to real time quantitative RT-PCR, and DUOX2 expression relative to β-actin was determined; error bars represent standard deviations; data are from triplicate samples; \*\*\**P* < 0.001 for: DMSO treated vs. IL-4 + IL-17A; IL-4 + IL-17A vs. Dex + IL-4 + IL-17A; and Dex + IL-4 + IL-17A vs. RU-486 + Dex + Il-4 + IL-17A. In these experiments, the effect of DMSO was studied because Dex and RU-486 were dissolved in DMSO. Middle panel shows a Western blot (50 µg lysate) from AsPC-1 cells in serum free medium treated with solvent (lane 1), IL-4 (50 ng/ml) plus IL-17A (50 ng/ml) plus solvent for 24 h (lane 2), or both cytokines alone (lane 3), IL-4 plus IL-17A plus Dex (lane 4), or both cytokines plus Dex plus RU-486 (lane 5). Pretreatment with Dex and/or RU-486 was for 30 min. The right panel shows the effect of Dex on IL-4 and/or IL-17A enhancement of MMP7 expression; \*\*\**P* < 0.001 for the effect of Dex on cytokine-enhanced MMP7 mRNA expression. **D,** Dex inhibits cytokine-induced DUOX protein expression and DNA damage response in CFPAC-1 cells. Western analysis of CFPAC-1 cells treated with LPS (1 μg/ml) alone, IFN-γ (25 ng/ml) alone, or both for 48 h with or without Dex pretreatment for 30 min. **E**, Flavin dehydrogenase and NADPH oxidase inhibitor diphenylene iodonium (DPI) exposure or silencing DUOX2 expression decreases IFN-γ-mediated extracellular H_2_O_2_ production in BxPC-3 cells. Control or DUOX2-specific siRNAs were transiently transfected into BxPC-3 cells; 24 h after transfection, BxPC-3 cells grown in serum-free medium were treated with 25 ng/ml of IFN-γ for another 24 h with or without DPI (1 µM). Tumor cells were then harvested, and extracellular H_2_O_2_ formation was measured for 1 h in the presence of ionomycin (1μM). Error bars represent standard deviations from triplicate experiments and 12 readings. *** *P*< 0.001 between the two conditions that have been compared. **F**, Dex suppresses cytokine-induced DUOX2 mRNA expression and H_2_O_2_ production in CFPAC-1 human pancreatic cancer cells. Left panel demonstrates real time PCR of CFPAC-1 cells grown in serum free medium and treated with either LPS (1 µg/ml) plus IFN-γ (25 ng/ml) with or without 1 µM Dex or IL-17A (50 ng/ml) + IFN-γ (25ng/ml) with or without 1 µM of Dex. Right panel demonstrates extracellular H_2_O_2_ production in CFPAC-1 cells treated with IL-17A (50 ng/ml) plus IFN-γ (25 ng/ml) with or without 1 µM of Dex. The enzymatic reaction systems contained ionomycin (1 μM). \*\*\**P* < 0.001 for cytokine-treated vs. solvent-treated cells and for Dex-treated, cytokine-exposed vs. cytokine-exposed tumor cells alone. All experiments were performed in triplicate.

RNA interference experiments (**Fig. 3B**) provided further support for the role of DUOX in enhancing IFN-γ-mediated Hif-1α expression and DNA damage. Control and DUOX1/2 specific siRNAs were transiently transfected into BxPC-3 cells; at 24 h following transfection, cells were cultured in serum-free medium with IFN-γ (25 ng/ml) for another 12 h. Cells were then collected and whole cell lysates were subjected to Western blot analysis. In the presence of control siRNA (**Fig. 3B**, left panel, lane 2), IFN-γ exposure activated STAT1 and S6, was associated with increased DUOX, Hif-1α, and JunB expression, and produced DNA double strand scission as indicated by enhanced γH2AX formation. In contrast, silencing DUOX1/2 with a specific siRNA abrogated IFN-γ-enhanced DUOX protein expression as well as IFN-γ-stimulated Hif-1α and JunB expression and the DNA damage response (compare lanes 2 and 4, **Fig. 3B**, left panel). In the right panel of **Fig. 3B**, we transfected CFPAC-1 cells with either a control siRNA or two different human DUOX2 siRNAs. 24 h following transfection the cells were treated with solvent (lane 1) or LPS (1 µg/ml) plus IFN-γ (25 ng/ml) for another 24 h; cell lysates were prepared for Western blotting using antibodies to the indicated proteins. We found that DUOX expression in CFPAC-1 cells was decreased by two different siRNAs. Diminished DUOX levels were accompanied by decreased expression of Hif-1α.

To extend our findings to other human pancreatic cancer cell lines, AsPC-1 cells were used to evaluate the potential roles of Dex and RU-486 on cytokine-mediated DUOX2 expression and DNA damage. As demonstrated in the left panel of **Fig. 3C**, although the DMSO solvent had no effect on the enhanced DUOX2 mRNA expression resulting from exposure to IL-4 (50 ng/ml) plus IL-17A (50 ng/ml), *P* < 0.001, 1 µM Dex significantly blunted cytokine-enhanced DUOX2 mRNA expression (\*\*\**P* < 0.001 for IL-4 plus IL-17A-treated cells vs. cells exposed to Dex plus cytokines). On the other hand, 1 µM RU-486 blocked the Dex-mediated decrease in DUOX2 expression (*P* < 0.001 vs. IL-4 plus IL-17A treatment with Dex and not RU-486). The Western blot shown in the middle panel of **Fig. 3C** supports the real time PCR results. IL-4 plus IL-17A increased DUOX protein expression in AsPC-1 cells with a concomitant increase in γH2AX signal; Dex decreased both DUOX and γH2AX expression levels in cytokine-treated cells (lane 4), whereas RU-486 pretreatment diminished the Dex-mediated decrease for both DUOX and γH2AX (lane 5). Although neither Dex nor RU-486 inhibited the effect of IL-4 on STAT6 phosphorylation, Dex activated, and RU486 inhibited, GR Ser 211 phosphorylation. The data shown in the right panel of **Fig. 3C** extends the effects of Dex to inhibition of cytokine-mediated upregulation of MMP7, a metalloproteinase important for the integrity of the extracellular matrix. In CFPAC-1 cells, as shown in **Fig. 3D**, either LPS (1 µg/ml; lane 2) or IFN-γ (25 ng/ml; lane 4) alone or in combination (lane 6) increased DUOX expression and DNA damage. Dex inhibited cytokine-mediated DUOX expression and DNA damage in CFPAC-1cells, as it did for the BxPC-3 and AsPC-1 cell lines. Dex treatment induced GR_S211_ phosphorylation but had no effect on total GR protein expression for the 24 h of its exposure.

Finally, to determine whether IFN-γ-induced DUOX2 and DUOXA2 expression was correlated with the functional activity of DUOX2, we examined extracellular H_2_O_2_ generation by BxPC-3 cells exposed to IFN-γ using the Amplex Red^®^ assay. When the Amplex Red reaction system contained the calcium ionophore ionomycin (1 µM), BxPC-3 cells that had been treated with IFN-γ produced significantly more H_2_O_2_ compared to cells that had not been incubated with IFN-γ (*P* < 0.001 vs. solvent treated, **Fig. 3E**). In this system, DPI (1 µM) decreased DUOX2-mediated H_2_O_2_ release by the IFN-γ-stimulated tumor cells to near-basal levels (*P* < 0.001 vs. IFN-γ-treated cells without inhibitor). Silencing DUOX2 with DUOX2-specific siRNA almost completely blunted IFN-γ-induced H_2_O_2_ production (*P* < 0.001 vs. control siRNA-treated cells exposed to IFN-γ). This data suggests that inhibiting the enzymatic activity of DUOX2 with the NOX/dehydrogenase inhibitor DPI or decreasing DUOX2 expression using RNA interference attenuates DUOX2-derived H_2_O_2_ production in BxPC-3 cells.

To corroborate further our observation that Dex-mediated decreases in cytokine-enhanced DUOX2 expression is associated with decreased H_2_O_2_ production in human pancreatic cancer cells, we also studied the CFPAC-1 cell line propagated in serum free medium and treated with DMSO as a solvent or Dex (1 µM) for 30 min followed by a 24 h exposure to LPS or IL-17A plus IFN-γ. DUOX2 mRNA expression and H_2_O_2_ formation were investigated in CFPAC-1 cells under these conditions. As demonstrated in the left panel of **Fig. 3F**, co-treatment with either LPS plus IFN-γ, or IL-17A plus IFN-γ produced a significant increase in DUOX2 RNA expression; Dex significantly inhibited the enhanced DUOX2 expression generated by both cytokine combinations. In the right panel, Amplex Red^®^ assay revealed that Dex significantly decreased IL-17A plus IFN-γ-related extracellular H_2_O_2_ generation in CFPAC-1 cells (*P* < 0.001 vs. IL-17A plus LPS without Dex).

### Nuclei from Dex-treated human pancreatic cancer cells contain activated GR and the necessary transcription corepressors to mediate the transcriptional repression of *DUOX2* gene induction by many cytokines

Two major mechanisms have been proposed to explain glucocorticoid-mediated repression of certain pro-inflammatory genes that are activated during the inflammatory response. For some genes (such as IL-6, collagenase, and NOSII) which lack functional GRE cis-elements in their promoter regions, but instead contain binding sites for other DNA-bound regulators, such as NF-κB and AP-1, ligand-activated GR can trans-repress gene expression by interfering with the function of these stimulatory transcription factors (39–41). On the other hand, for such genes as thymic stromal lymphopoietin (TSLP), glucocorticoids induce direct trans-repression via ligand-activated GR binding to negative GC response elements (IRnGRE). These elements act on agonist-liganded GR, promoting the assembly of cis-repressing complexes, containing liganded-GR-NCoR1/NCoRII, and histone deacetylases (35).

Thus, a possible mechanism for the Dex-mediated decrease in cytokine-related DUOX2 expression may be that the activated GR directly binds to the DUOX2 promoter, recruiting various co-repressors to repress IFN-γ-enhanced DUOX2 expression. To explore this possibility, the initial 6-kilobase segments of the human DUOX2 promoter were scanned to identify potential activated GR binding sites matching the canonical IRnGRE binding motif 5′-<u>CTCC</u>N<u>GGAG</u>-3′ (35), in which the underlined letters correspond to conserved palindromes and N denotes any of the 4 nucleotides. We identified such a site in the human DUOX2 promoter region with the sequence 5’-CAG<u>CTCC</u>A<u>GGAG</u>TTC-3’, which is a match for the reported GR binding sequence. We then wanted to understand whether activated GR and other co-repressors exist in human pancreatic cancer cells. The nuclear distribution of various transcription factors was evaluated following exposure to IFN-γ (25 ng/ml, 3 h) with or without a 30 min pretreatment with the inhibitors Dex or RU-486. As shown in **Fig. 4A** for BxPC-3 cells and **Fig. 4B** for ASPC-1 cells, GR, co-repressor NCoR1/2, and HDACs1-3 constitutively exist in the nuclei of these tumor cells; IFN-γ, Dex, and RU-486 have no effect on the relative nuclear distribution of the components of the cis-repressing complex, except for GR phosphorylated at Ser211 for which Dex enhanced, and RU-486 inhibited, nuclear translocation. Moreover, as shown in the right panel of **Fig. 2A**, Dex had no effect on IFN-γ-induced STAT1 nuclear distribution in BxPC-3 cells. Supplementary Fig. S5 also demonstrates that CFPAC-1 cells constitutively express nuclear GR, NCoR1/2 and various HDACs; neither Dex nor RU-486 treatment influenced the nuclear distribution of these factors. Only the nuclear translocation of GR phosphorylated at Ser211 was enhanced by Dex and inhibited by RU-486, similar to our observation using BxPC-3 cells. In summary, the DUOX2 promoter contains canonical negative GRE elements, and Dex enhances Ser211 activated GR translocation into the nucleus in human pancreatic tumor cells constitutively expressing various co-repressors in their nuclei. Thus, the conditions exist in PDAC cells to permit activated GR to bind directly to the human DUOX2 promoter which would permit the assembly of a repressor complex capable of inhibiting IFN-γ-enhanced DUOX2 expression.

**Figure 4.**
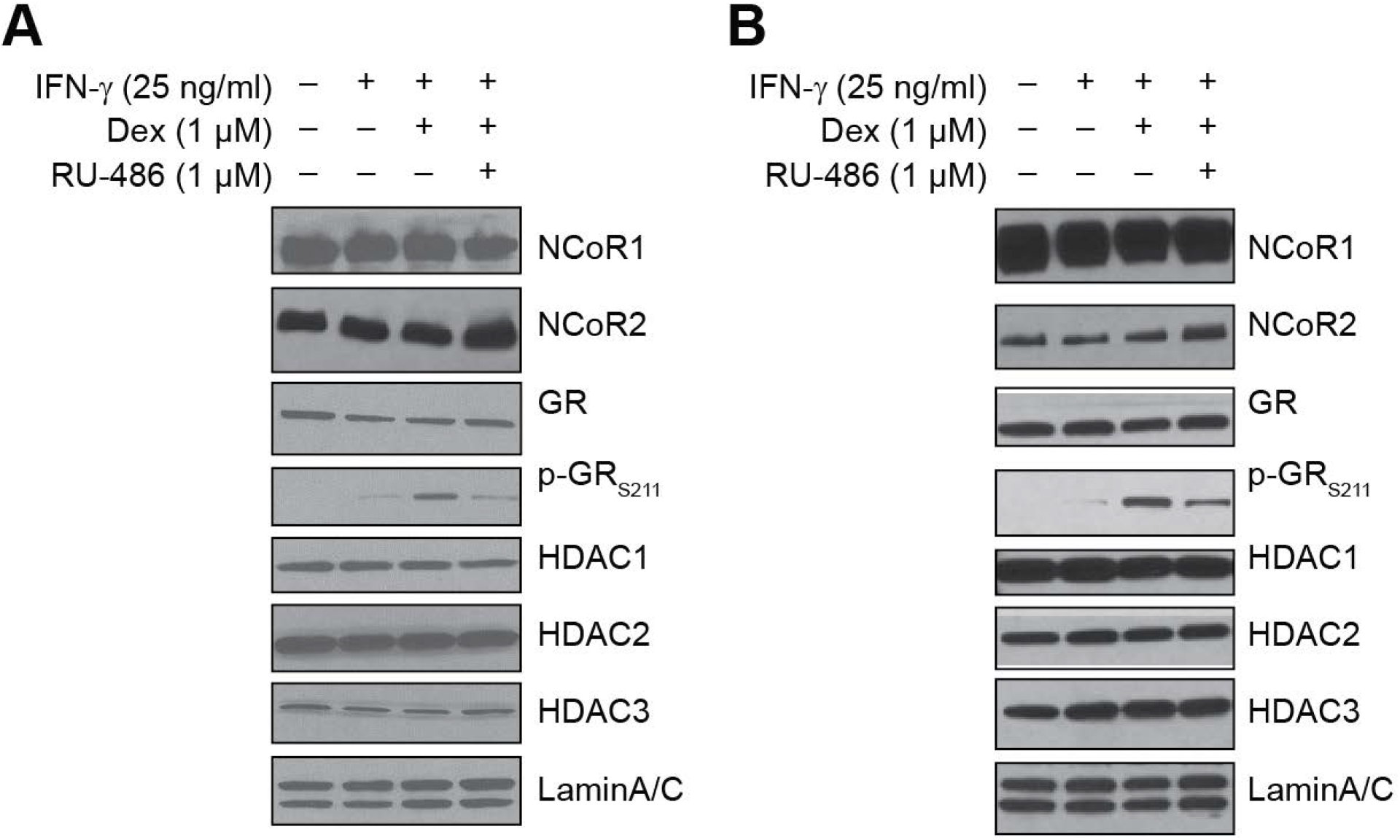
Human DUOX2 promoter contains a putative Negative Glucocorticoid Receptor Binding Element (nGRE: <u>CTCCaGGAG</u>). Dex-stimulated human pancreatic cancer cell nuclei contain activated GR and all the necessary transcriptional corepressors that could participate in the transcriptional repression by Dex of cytokine-enhanced DUOX2 gene expression. **A,** BxPC-3 cells and **B,** AsPC-1cells grown in serum free medium were treated with IFN-γ (25 ng/ml), Dex (1 µM) or RU-486 (1 µM) for 3 h; 20 μg of nuclear extract was subjected to Western analysis to determine target protein expression. Lamin A/C was used as a loading control for nuclear protein. The data are representative of experiments performed in triplicate.

### Dex does not alter human pancreatic cancer cell proliferation *in vitro* but suppresses the growth of BxPC-3 tumor xenografts and the expression of DUOX2, VEGF-A, and inflammatory cell infiltrates *in vivo*

Because pro-inflammatory cytokines enhance ROS production and Hif-1α expression that is significantly decreased by Dex pre-treatment, and because our studies have demonstrated that IL-4 exposure enhances DUOX2 expression in BxPC-3 cells (**Fig. 1D**) and significantly increases NOX-dependent growth of colon cancer cells *in vitro* (42), we evaluated the direct effect of Dex on PDAC tumor cell proliferation. Using the MTT assay, we measured the effect of Dex on cell growth in BxPC-3 and MIA-PaCa cells. As shown in **Fig. 5A**, there was no significant difference in growth rates when solvent- and Dex-treated cells of either cell line were compared over three days of observation. Our data are consistent with those of Rhim et al. showing that Dex treatment *in vitro* of PanIN-derived epithelial cells had no effect on their morphology, proliferation rate, or the expression of epithelial or mesenchymal markers (3). However, when mice bearing BxPC-3 xenografts were treated with two different dose levels of Dex (6.25 or 12.5 mg/kg subcutaneously every 12 h for 22 doses beginning at the time the tumors reached a volume of ≈150 mm^3^), growth of the xenografts was significantly decreased by both Dex treatment regimens (**Fig. 5B**). On the other hand, for MIA-PaCa xenografts that do not demonstrate DUOX2 upregulation following cytokine exposure *in vitro*, Dex treatment had no effect on xenograft growth (**Fig. 5B**).

**Figure 5.**
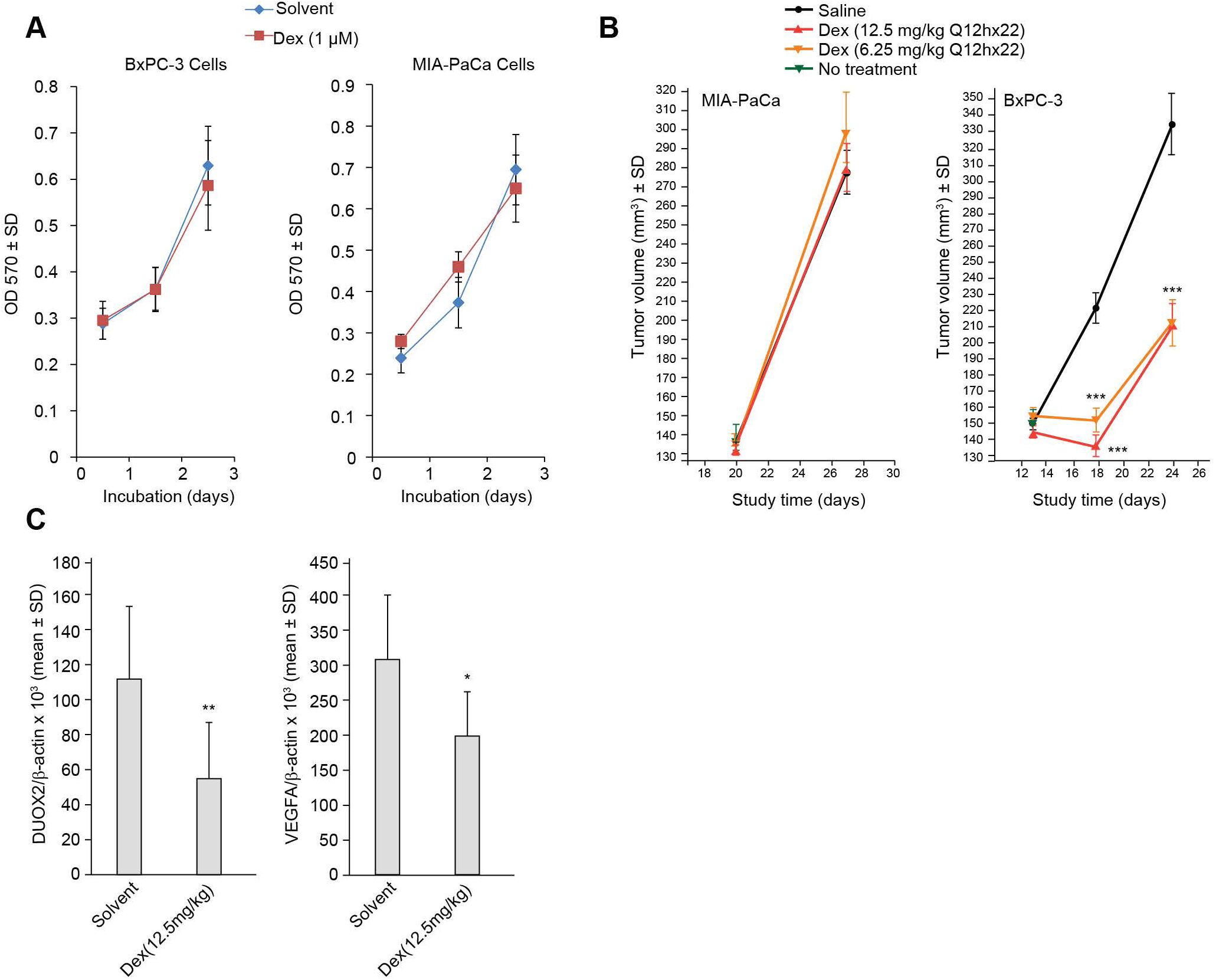
Dexamethasone has no effect on human pancreatic cancer cell proliferation *in vitro* but suppresses the growth of BxPC-3 xenografts and significantly decreases the expression of VEGF-A and inflammatory cell infiltrates *in vivo* in athymic mice. **A,** Dex does not alter human pancreatic cancer cell proliferation *in vitro*. BxPC-3 and MIA-PaCa cells grown in complete medium were treated with DMSO or 1 µM Dex for 3 days. MTT assays were then performed to evaluate cell proliferation. Data are expressed as means ± SD for triplicate experiments. **B,** Effect of Dex on human pancreatic cancer cell growth *in vivo*. When xenografts reached 150 mm^3^, treatment was begun with saline or two different dose levels of Dex (6.25 and 12.5 mg/kg) administered subcutaneously every 12 hours for 22 doses. Mice were observed for tumor growth. Dex administration produced no effect on MIA-PaCa xenograft growth; both Dex dose schedules significantly decreased tumor size at study days 18 and 24, \*\*\**P* < 0.001 for the comparison of saline-treated with Dex-treated animals carrying BxPC-3 xenografts. **C,** Dex inhibits DUOX2 and VEGF-A mRNA expression in BxPC-3 pancreatic cancer xenografts. Additional groups of 8 mice in each treatment program were sacrificed 4 h after the final dose of 12. 5 mg/kg Dex for gene expression analysis. Tumors were resected, flash frozen, and stored at -80°C until processing. Quantitative RT-PCR results are demonstrated for DUOX2 and VEGF-A mRNA expression from BxPC-3 xenografts treated with either Dex or saline. \*\**P* < 0.01 and \**P* = 0.01 for the comparison of DUOX2 and VEGF-A expression, respectively, in BxPC-3 xenograft tumors treated with Dex vs. vehicle.

To evaluate potential mechanisms that might explain the effect of Dex on BxPC-3 tumor growth *in vivo*, we examined the influence of the glucocorticoid on the expression of DUOX2 and VEGF-A. As demonstrated in **Fig. 5C**, Dex treatment significantly decreased mRNA expression for both DUOX2 and VEGF-A in xenografts removed four hours after the last dose of glucocorticoid.

Finally, as has been demonstrated in other tumor xenograft models (43), we found that Dex treatment significantly decreased the infiltration of leukocytes (including neutrophils) and macrophages into BxPC-3 xenografts (**Fig. 6 and Fig. 7**). CD3^+^ and CD8^+^ T cells as well as B cells in the tumor microenvironment were also significantly diminished. Decreased immune and inflammatory cell infiltration in this case was accompanied by a significant increase in tumor cell apoptosis (measured as activated caspase 3) and a significant decrease in tumor cell proliferative index (Ki-67) consistent with the Dex-related inhibition of BxPC-3 xenograft growth (**Fig. 5**).

**Figure 6.**
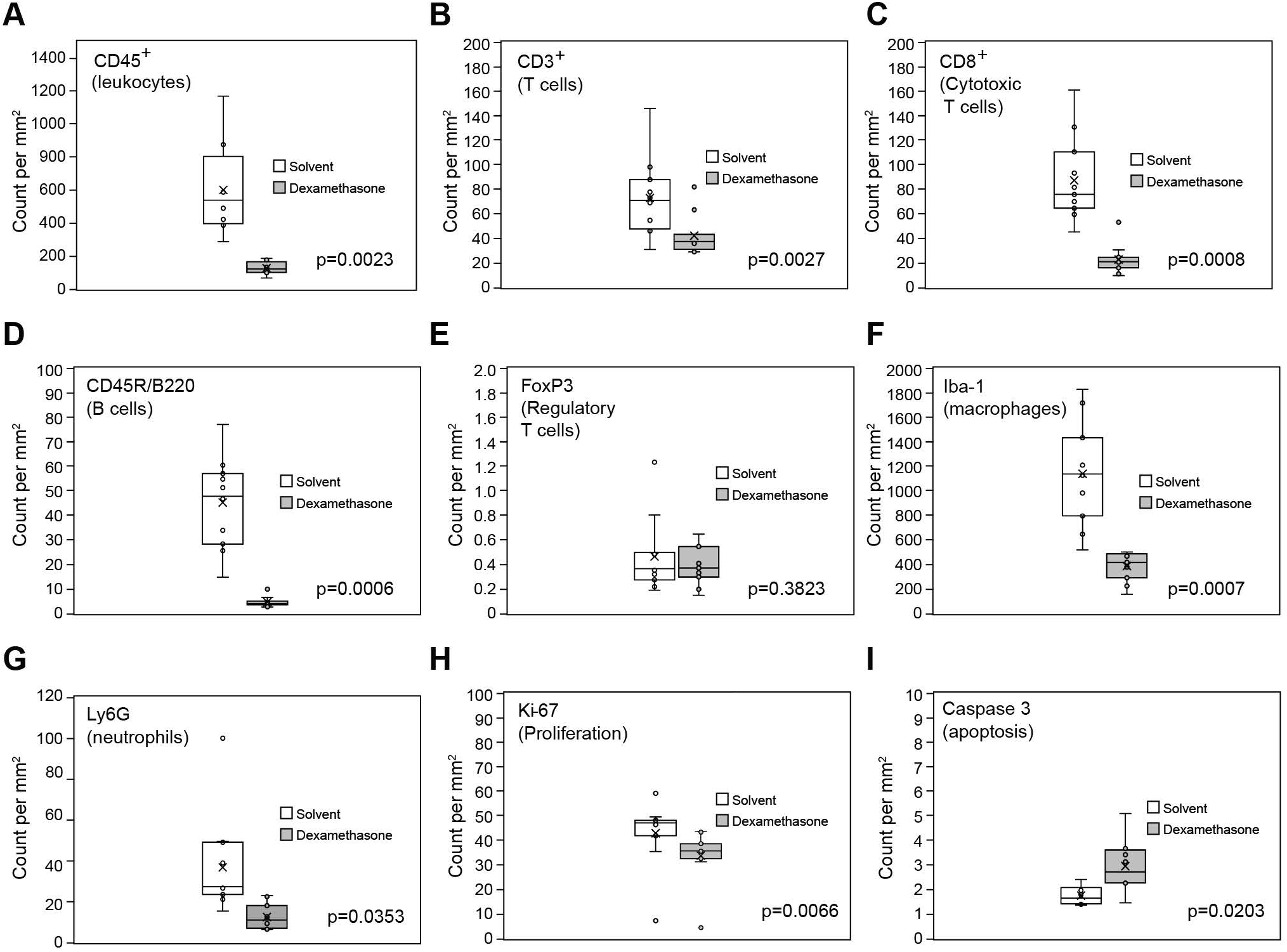
Immunohistochemical analysis of inflammatory and immune cell numbers, and proliferation markers in BxPC-3 xenografts. Paraffin-embedded tumor samples from 8 solvent-treated mice (open box plots) and 8 dexamethasone-treated mice (filled box plots) were subjected to IHC analysis using antibodies for the indicated markers of each inflammatory or immune cell type in the xenografts (**A-G**). Proliferation and apoptosis markers for the tumor cells were assessed using anti-Ki-67 and anti-caspase 3 antibodies, respectively (**H, I**). The ‘x’ and horizontal line in each box represent the mean and median values, respectively. Evaluation of the significance of each group comparison between solvent-treated and Dex-treated mice is provided for the individual panels.

**Figure 7.**
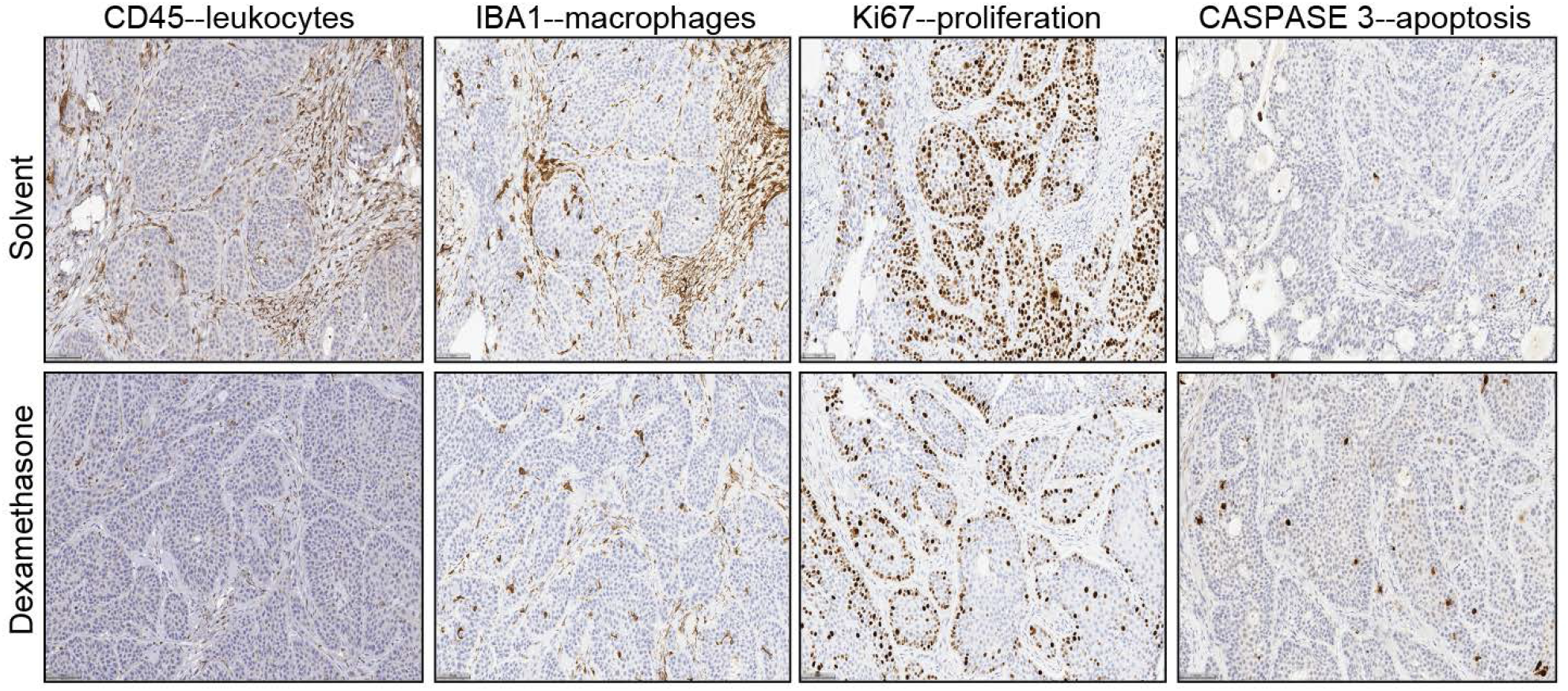
Representative images from the IHC analysis of BxPC-3 tumor samples are shown for both solvent-and dexamethasone-treated mice. The dense infiltration of leukocytes and macrophages into the interstitial microenvironment of the tumor and the significant reduction in inflammatory cell numbers following dexamethasone treatment is demonstrated in the panels on the left. On the other hand, a significant decrease in proliferation (measured by Ki67 staining) and increase in apoptosis (caspase 3) is shown predominantly for tumor cells rather than microenvironmental infiltrates in the two panels on the right of the figure. A scale bar (100 µm) is shown at the lower left margin of each image.

## Discussion

In these experiments, we sought to provide further evidence supporting a role for DUOX2 in the pathophysiological behavior of PDACs. Our laboratory has established that DUOX2 is highly expressed at the protein level in patients with chronic pancreatitis (18), as well as pancreatic intraepithelial neoplasia and early stages of PDAC compared to tumor-adjacent, histologically normal pancreas (19), and that the survival of patients with PDAC is significantly diminished in the presence of high level DUOX2 mRNA expression (21). We have also demonstrated that several cytokines known to play an important role in tumor-associated inflammation upregulate DUOX2 expression in PDAC cells (21). Because Dex significantly decreased the growth rate of xenografts developed from the AsPC-1 human PDAC cell line (44) and a patient-derived xenograft (PDX) model (28), as well as post-surgical recurrence and invasiveness in other PDAC animal models (45), we examined whether Dex altered DUOX2 expression in human PDAC cells and xenografts. *In vitro,* Dex significantly inhibited the increase in DUOX2 expression produced by a wide variety of pro-inflammatory molecules (including IFN-γ, LPS, IL-4, IL-17A, and IFN-β); Dex administration *in vivo* significantly decreased the growth of BxPC-3 xenografts. The studies presented herein have attempted to characterize further the mechanisms by which Dex alters cytokine-mediated DUOX2 expression and function.

Dex interfered with the upregulation of DUOX2 expression by pro-inflammatory cytokines in several human PDAC cell lines, and this GC-mediated effect appeared to be associated with GR activation. In previous studies, we have shown that increased DUOX2 levels following exposure of PDAC cells to LPS or IL-17A may be mediated by translocation of p65 to the tumor cell nucleus and binding to the DUOX2 promoter (18, 21). Thus, the effects of Dex on DUOX2 expression that we observed *in vitro* for these cytokines could be due to trans-repression of NF-κB binding to DNA by activated GR (46). In a similar fashion, Dex-related interference with the function of AP-1 might diminish cytokine-related activation of multiple downstream signaling pathways (40) that contribute to a pro-angiogenic milieu in PDACs (19).

However, although trans-repression by Dex may help to explain its effects on LPS- or IL-17A-related upregulation of DUOX2, this effect is unlikely to be the complete explanation for our results with IFN-γ. Despite the observation that LPS alone or in combination with IFN-γ can enhance p65 translocation in BxPC-3 cells, IFN-γ alone was ineffective in this regard [see Fig. 4F in (18)]. The MEK inhibitor U0126 can decrease c-Jun and JunB nuclear expression in BxPC-3 cells [see Fig. 7F in (19)], but it had no effect on IFN-γ-induced DUOX2 expression in the current work (Supplementary Fig. S4, middle panel). Thus, trans-repression of AP-1 or NF-κB transcriptional activity may not play singular roles in Dex-mediated inhibition of IFN-γ-related upregulation of DUOX2.

By directly suppressing STAT1 mRNA and protein expression, GCs can inhibit IFN-γ signaling and the expression of downstream genes, such as IRF-1, in PBMCs (47). However, in BxPC-3 cells, as shown in the right panel of **Fig. 2A**, Dex had no effect on IFN-γ-induced STAT1 protein expression. Furthermore, although IFN-γ can induce STAT1 phosphorylation at both Y701 and S727 and expression of its downstream target IRF-1, neither Dex nor RU486 altered IFN-γ-enhanced STAT1 activation or IRF-1expression (**Fig. 3A**, compare lane 3 with lane 4, left panel). These data suggest that direct inhibition of JAK-STAT signaling is not solely responsible for the effect of Dex on IFN-γ-enhanced DUOX2 expression in BxPC-3 cells.

Thus, it seems possible, at least for our studies with IFN-γ, that other mechanisms, including the direct binding of activated GR to the DUOX2 promoter, with the subsequent recruitment of various corepressors, could contribute to the repression of IFN-γ-enhanced DUOX2 expression. We have shown that all necessary conditions exist in the nuclei of several PDAC cell lines, including potential activated GR binding sites matching the canonical IRnGRE binding motif, to permit GR-mediated inhibition of DUOX2 expression. However, despite performing chromatin immunoprecipitation and DNA affinity precipitation studies coupled with Western analysis to evaluate potential binding of ligand-activated GR to possible GR binding sites in the human Duox2 promoter (data not shown), we were unable to develop evidence to support this mechanism. Hence, the role of transcriptional co-repression in explaining our results for IFN-γ remains hypothetical.

To evaluate the possible clinical relevance of our observation that Dex significantly decreased cytokine-enhanced DUOX2 expression in cell culture, we examined the effect of Dex treatment on tumor growth, gene expression, and inflammatory cell infiltrates in PDAC xenografts. In a previous study, we found that DUOX2 expression at the mRNA level was dramatically upregulated, absent any therapeutic intervention, during a single passage of BxPC-3 cells as xenografts in athymic nude mice (18). In the current studies, we found that treatment with Dex significantly decreased the expression of DUOX2 and VEGF-A mRNA in our animal model (**Fig. 5C**) while, at the same time, significantly decreasing xenograft development (**Fig. 5B**) and the expression of the Ki-67 proliferation marker (**Fig. 6H and Fig. 7**). Tumor cell growth *in vitro* was not affected by treatment with Dex (**Fig. 5A**). These results suggest that the loss of a pro-angiogenic oxidative milieu could play an important role in fostering a pro-apoptotic tumor microenvironment (**Fig. 6I and Fig. 7**). For the MIA-PaCa line, which does not upregulate DUOX2 *in vitro* or following passage as a xenograft (18), Dex produced no effect on tumor cell proliferation or xenograft growth (**Fig. 5**).

In view of the pleiotropic effects of Dex on the inflammatory/immune response (25), there are several potential explanations for our *in vivo* observations beyond an effect on the DUOX2 promoter. Although conflicting reports exist (48), recent studies support the observation that Dex produces little, if any, change in the proliferation rate of PDAC cell lines *in vitro* (29,44,45). These reports support the hypothesis that the growth inhibition we found *in vivo* may be related to one or more effects of Dex on the interaction between BxPC-3 xenografts and their ecosystem, interactions that could occur even within the immunocompromised context of the athymic nude mice used for our experiments (49). In particular, the significantly decreased expression of tumoral DUOX2 that we observed for Dex-treated mice bearing BxPC-3 xenografts, in conjunction with decreased VEGF-A expression, may lead to lower levels of H_2_O_2_ formation and related DNA damage and Hif-1α-mediated angiogenesis in both the tumor and microenvironment; these factors could contribute to impaired xenograft growth. Dex is also known to decrease LPS- and IFN-γ-enhanced transcription of the components of the NOX2 complex in macrophage-like cells (50), leading to decreased oxidation of extracellular matrix proteins (51). These results are consistent with our observation that Dex significantly diminished the cytokine-enhanced expression of MMP7 in BxPC-3 cells (**Fig. 3C**); if a similar phenomenon occurs *in vivo*, it could also contribute to the inhibition of xenograft growth we observed.

Finally, we found that BxPC-3 xenografts from Dex-treated mice demonstrated a significant decrement in the extent of peri-tumoral inflammatory and immune cell infiltrates (**Fig. 6A-G**). Dex-related reduction in neutrophil and macrophage penetration of the interstitial space of the xenografts would likely diminish the secretion of pro-inflammatory cytokines capable of upregulating DUOX2 expression and the subsequent generation of a reactive oxygen cascade. At the same time, the Dex-related loss of CD8^+^ T cell numbers that we observed would have been unlikely to substantively exacerbate the well-known limitations in functional immune and proliferative capacity of T cells derived from athymic nude mice hosts (52, 53).

In summary, our studies suggest that the anti-invasive and anti-proliferative effects of Dex that have been observed previously in both xenograft and genetically-engineered models of PDAC (2, 44) could be mediated, in part, by inhibition of a DUOX2-related cascade of pro-inflammatory ROS, thereby significantly reducing the local production of pro-angiogenic factors that play an essential role in pancreatic tumor cell growth.

## 2 Abbreviations

DUOX2: dual oxidase 2
VEGF: vascular endothelial growth factor
ROS: reactive oxygen species
PDAC: pancreatic ductal adenocarcinoma
PanIn: pancreatic intraepithelial neoplasia
GR: glucocorticoid receptor
HDAC: histone deacetylase
NOX: NADPH oxidase
Dex: dexamethasone
DPI: diphenylene iodonium
NAC: n-acetyl-L-cysteine

## Footnotes

1 **Financial Support** This project has been funded in whole or in part with federal funds from the National Cancer Institute, National Institutes of Health (ZIA BC011076-05), and under Contract No. HHSN261200800001E. The content of this publication does not necessarily reflect the views or policies of the Department of Health and Human Services, nor does mention of trade names, commercial products, or organizations imply endorsement by the U.S. Government.

## Notes

**Disclosure of Potential Conflicts of Interest**: The authors declare no potential conflicts of interest.

### Competing Interest Statement

The authors have declared no competing interest.

