## Supplemental Figure Legends and Figures for "Dexamethasone Inhibits Cytokine-Induced, DUOX2-Related VEGF-A Expression and DNA damage in Human Pancreatic Cancer Cells and Growth of Pancreatic Cancer Xenografts"

**Supplementary Figure S1.**

Comparison of the effects of IFN-γ and Dex on DUOX2 and VEGF-A induction in two human pancreatic cancer cell lines. In the left panel, BxPC-3 or MIA-PaCa cells grown in serum free media were treated with solvent, IFN-γ (25 ng/ml) alone, or IFN-γ plus Dex for 24 h. Cells were collected, and RNA was extracted for quantitative PCR analysis of DUOX2 and VEGF-A mRNA expression relative to β-actin. ***$P < 0.001$ for solvent vs. cells treated with IFN-γ (25 ng/ml); ***$P < 0.001$ for cells treated with IFN-γ (25 ng/ml) for 24 h alone vs. Dex plus IFN-γ (25 ng/ml) for 24 h. For experiments shown in the right panels, MIA-PaCa human pancreatic cancer cells were propagated in serum free medium and then treated with solvent (lane1), Dex (lane 2), IFN-γ (lane 3), or IFN-γ plus Dex for 24 h; 50 µg of whole cell extract was used for Western analysis using specific antibodies as indicated in the figure. The data are representative of triplicate experiments.

**Supplementary Figure S2.**

Flavin dehydrogenase and NOX inhibitor DPI and the reduced thiol NAC attenuate IFN-γ-enhanced VEGF-A expression in BxPC-3 cells. BxPC-3 cells in serum free medium were pretreated with 1 µM DPI or 10 mM NAC for 30 min and then exposed to IFN-γ (25 ng/ml) for 24 h; quantitative PCR analysis was then performed to determine DUOX2 and VEGF-A mRNA expression relative to β-actin. For either VEGF-A or DUOX2, ***$P < 0.001$ for the comparison of solvent vs. cells treated with IFN-γ (25 ng/ml) alone; VEGF-A, ***$P < 0.001$ for cells treated with IFN-γ (25 ng/ml) for 24 h alone vs. IFN-γ plus either DPI or NAC for 24 h; neither DPI nor NAC inhibited IFN-γ-enhanced expression of DUOX2. Results determined from at least three identical experiments.

**Supplementary Figure S3.**

Silencing DUOXA2 decreases IFN-γ-induced DUOXA2 and VEGF-A expression in BxPC-3 cells. Control or DUOXA2-specific siRNAs were transiently transfected into BxPC-3 cells. At 24 h after transfection, cells were incubated in serum-free medium with or without IFN-γ (25 ng/ml) for another 24 h. Relative DUOXA2 (left panel) and VEGF-A (right panel) mRNA expression levels were normalized to β-actin. ***$P$ < 0.001 for solvent vs. cells treated IFN-γ (25 ng/ml); ***$P$ < 0.001 for cells treated with IFN-γ (25 ng/ml) plus control siRNA for 24 h vs IFN-γ plus DUOXA2 siRNA. Data represent results from three experiments.

**Supplementary Figure S4.**

Real time PCR assay of BxPC-3 cells grown in serum free medium and treated with a MEK inhibitor U0126 (10 μM) or DPI (1μM) for 30 min followed by exposure to IFN-γ (25 ng/ml) for 24 h. Relative VEGF-A (left panel), DUOX2 (middle panel), and STAT1 (right panel) mRNA expression normalized to β-actin. ***$P$ < 0.001 for solvent vs. cells treated IFN-γ (25 ng/ml); ***$P$ < 0.001 for VEGF-A expression in cells treated with IFN-γ (25 ng/ml) alone vs. IFN-γ plus inhibitors. Neither U0126 or DPI addition inhibited DUOX2 or STAT1 expression that had been enhanced by IFN-γ. Data represent three experiments.

**Supplementary Figure S5.**

CFPAC-1 cells grown in serum free medium were treated with LPS (1 μg/ml), Dex (1 μM) or RU-486 (1 μM) as described in the figure, and 20μg of nuclear extract was subjected to Western analysis to determine target protein distribution in the nucleus using specific antibodies as indicated in the figure.  Lamin A/C was used as a loading control for nuclear protein.  Results are representative of three different experiments.

### Supplementary S1

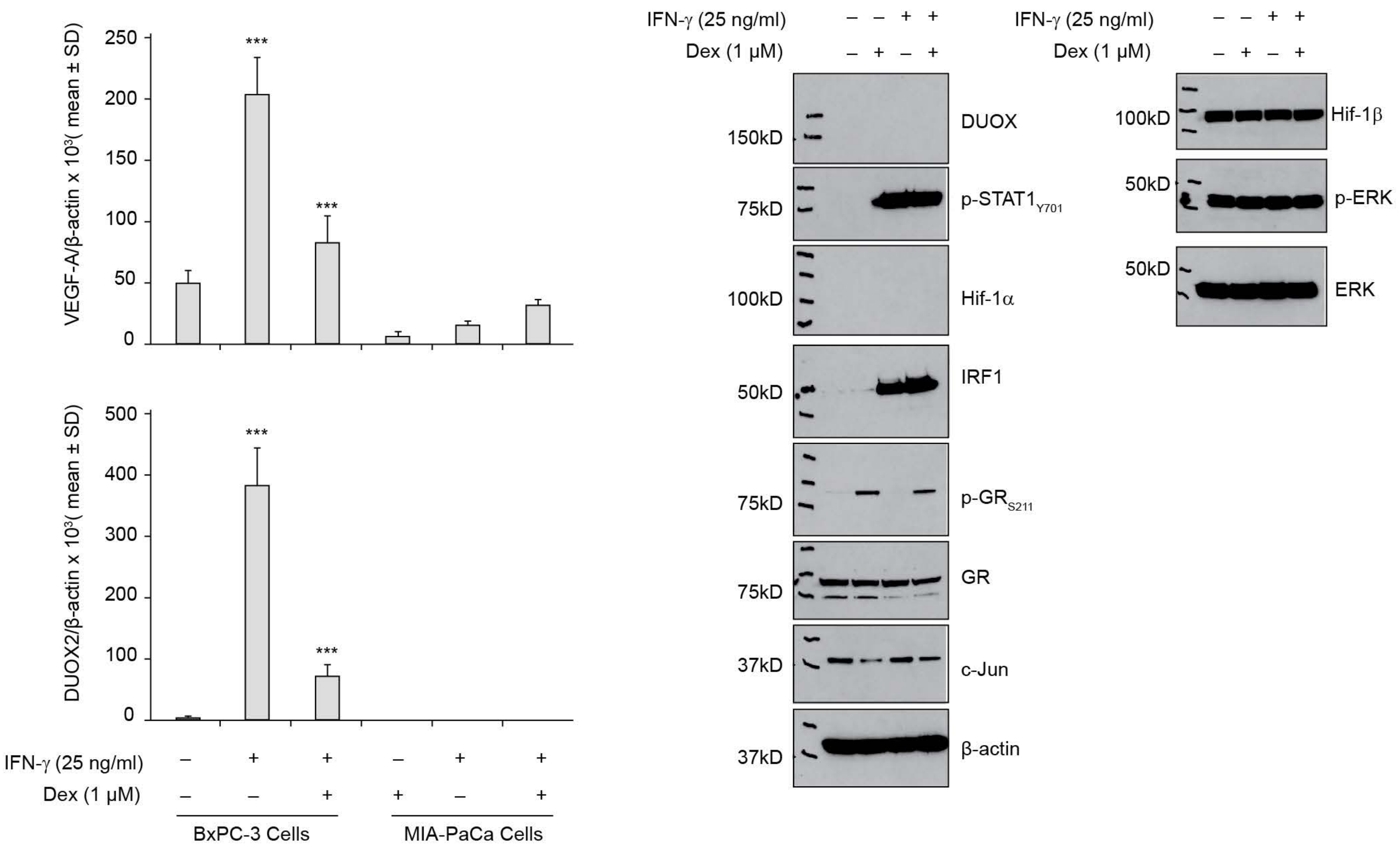

### Supplementary S2

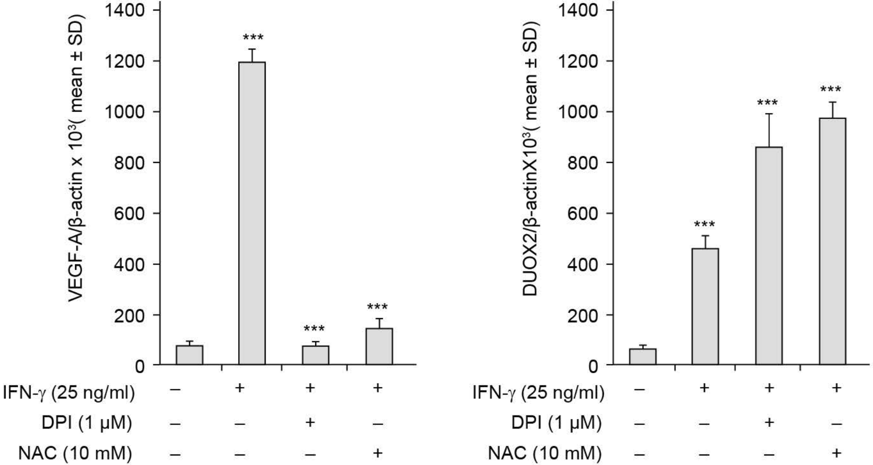

### Supplementary S3

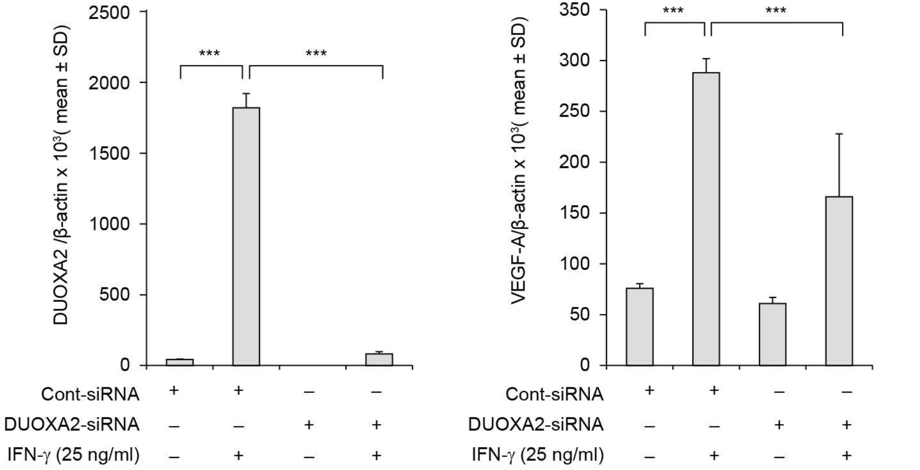

### Supplementary S4

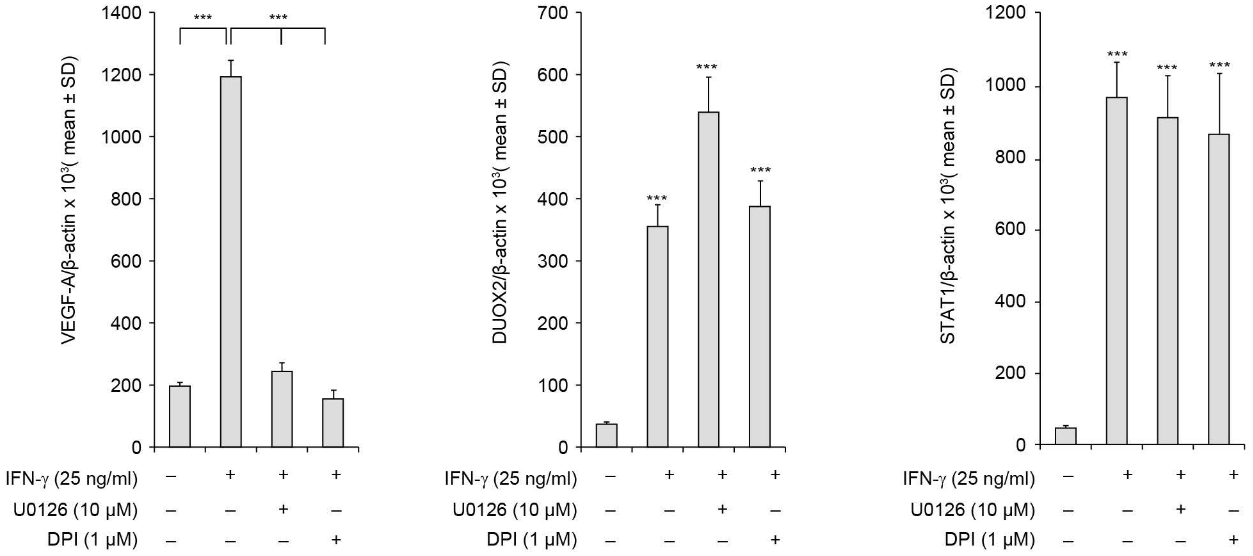

### Supplementary S5

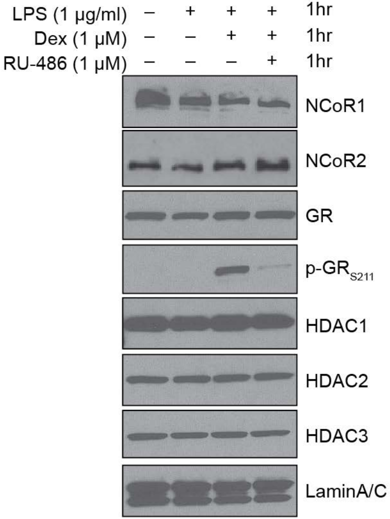
